# Appellian Neural Networks

**DOI:** 10.64898/2026.06.03.729998

**Authors:** Serhii Bahdasariants, Lauren Parola, Kriti Kacker, Ariel K. Feldman, Zachary Zdobinski, Inseung Kang, Douglas J. Weber

## Abstract

The movement of articulated bodies, whether biological or robotic, involves two interrelated computational challenges. Inverse dynamics determines the forces or torques required to achieve a specified movement, while forward dynamics computes the resulting accelerations from given forces or torques. Although both computations are governed by the same equations of motion, they are typically implemented as separate inverse and forward mappings in both model-based and data-driven approaches. This division can obscure the underlying structure common to both problems. Here, we introduce the Appellian Neural Network (ANNet), a physics-informed neural network that unifies both computations through a shared learned representation by modeling a single scalar quantity from classical mechanics, specifically Appell’s acceleration energy. Inverse dynamics is obtained by differentiating the learned energy function with respect to acceleration to recover joint torques. Forward dynamics is computed without retraining by embedding the same learned energy landscape into an optimization objective, where the unregularized minimum satisfies the Gibbs–Appell equation. The resulting accelerations are then integrated forward in time. ANNet is evaluated using a double pendulum paradigm. For trajectories not encountered during training, both inverse and forward simulations demonstrate accuracy and real-time capability. Both computations propagate input noise similarly to their analytical counterparts, and adaptation of the inverse dynamics to a modified pendulum generalizes to forward simulation without additional forward-specific training. These results establish a first-principles approach for employing a single learned representation to support both inverse and forward dynamic computations.

## Introduction

The ability of biological systems to generate a wide repertoire of precise and coordinated movements, from the rhythmic axial undulations of the lamprey^1–3^ to the intricate dexterity of the human hand^4,5^, is a hallmark of animal life. Underlying this capability is the computational challenge of managing the complex physics of movement^6,7^. To produce any goal-directed action, the nervous system must compute the complex, nonlinear relationship between the forces or torques generated by muscles and the resulting motion of the body’s articulated segments^8–11^. This process involves transforming a desired motor outcome into the specific pattern of muscle activations required to achieve it, a task complicated by the body’s intersegmental mechanics and its interaction with a variable environment^12,13^.

To solve the complex physics of movement, nervous systems employ two fundamental strategies, feedback and feedforward control^14,15^. Feedback control is a reactive process that uses continuous streams of sensory information primarily from vision^16–18^, touch^19^, and proprioception^20^ to detect deviations from an intended movement and generate real-time corrections. This mechanism is essential for maintaining accuracy and compensating for unexpected perturbations^21^. However, its effectiveness is limited by time delays inherent in sensory processing and neural transmission^22,23^, which constrain how rapidly the nervous system can generate a response. To reduce dependence on feedback-mediated corrections, organisms rely heavily on feedforward control, a predictive strategy that uses prior experience and internal knowledge of the body’s dynamics to generate motor commands in advance^24,25^. By anticipating the forces required to produce an intended action, feedforward control enables rapid, coordinated behaviors that would be impossible under feedback control alone, making it a key component of skilled movement^26^. Optimal motor performance is widely thought to arise from the integration of both predictive feedforward planning and reactive feedback correction^27,28^, enabling motor behaviors to be flexibly adapted to different objectives, such as minimizing metabolic cost or maximizing movement smoothness.

Central to the concept of feedforward control is the hypothesis that the brain constructs and maintains internal models, neural representations that mimic the input-output characteristics of the motor system and the environment^7,29,30^. Because internal models are inferred from behavioral signatures (adaptation, after-effects, anticipatory compensation) rather than observed directly, the term names a computational-level hypothesis about what the nervous system achieves rather than a commitment to how, or whether, those computations are realized as discrete neural objects. These models are generally categorized into two complementary types, forward and inverse models^31^. A forward model predicts the sensory consequences of a motor command. Given the current state of a limb and a set of motor commands, it simulates the causal flow of dynamics to predict the limb’s next state (e.g., its future position and velocity). This predictive capacity is crucial for overcoming sensory delays^22^, distinguishing self-generated sensations from external ones^32^, and providing a rapid estimate of the body’s state for online error correction^33^. Conversely, an inverse model performs the opposite transformation. It computes the motor commands required to achieve a desired sensory outcome or state transition^34,35^. By inverting the system’s dynamics, an inverse model functions as a controller, enabling the brain to generate the appropriate feedforward commands to execute a planned movement^36^.

An open question in motor neuroscience is whether these forward and inverse models represent separate computational processes, likely implemented in distinct neural circuits^31,37^, or whether they could arise from a single, unified neural substrate^38–40^. The prevailing view has treated them as separate, largely for mathematical reasons. Forward dynamics maps the current state and applied torques to the resulting acceleration 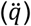, while inverse dynamics maps the current state and a desired acceleration to the required torques (*τ*)^41^. These are different input-output mappings that, in conventional machine learning, would necessitate different models or networks. This argument, however, concerns a representation rather than a nervous system, because the separation follows from the decision to describe dynamics as a map between states and torques, and need not survive a formulation in which both quantities are recovered from a common underlying object. Separate neural implementation is consistent with evidence of functional dissociations in neurological disorders and experimental findings suggesting distinct neural correlates for predictive and control-related signals^42–44^. However, arguments for a more unified architecture have been advanced, often on the grounds of efficiency^45,46^. Maintaining two separate, complex models for the same physical system could be metabolically and computationally costly, whereas a unified model could leverage shared representations to learn more efficiently. Such arguments are normative rather than evidential, because they identify what would be advantageous, not what is implemented. Recent findings suggest that some brain regions, such as the cerebellum and parietal cortex, are engaged during tasks thought to require both model types, leaving open the possibility of a shared substrate^47^. Co-engagement of this kind does not yet constrain the hypothesis space strongly. At the spatial and temporal resolution of current measurements, a genuinely shared computation and two interdigitated but functionally distinct circuits are difficult to tell apart, and different underlying organizations can support indistinguishable behavior.

Here we take the possibility of a shared substrate as motivation and ask a question that can be settled formally, namely whether the separation of the two computations is mathematically necessary. We demonstrate how a single neural network can function as a shared component of both a forward and an inverse dynamics model. Our approach is grounded in the Gibbs–Appell formulation of analytical mechanics and Gauss’s principle of least constraint, which reformulates system dynamics as an optimization problem. This perspective reveals that a single scalar function, the Appell acceleration energy, determines the equations of motion of a mechanical system once the applied generalized forces are specified. We demonstrate that a physics-informed ‘*Appellian’* neural network (ANNet) can be trained to approximate this scalar energy function. For inverse dynamics, the network learns by minimizing the error between the analytical derivative of its scalar output (with respect to acceleration) and the true joint torques, a process enabled by automatic differentiation (see Fig. 1). For forward dynamics, the same trained network is repurposed without any modification. By embedding the learned energy landscape into an optimization objective, the system solves for the accelerations that satisfy the physical principle of least constraint at each time step. Our central hypothesis is that a neural network trained to solve inverse dynamics by learning the scalar Appell acceleration energy encodes the inertial and Coriolis terms necessary and sufficient to solve forward dynamics when combined with a model of the applied generalized forces (e.g., gravity). We test this hypothesis using a simulated double pendulum, a model system that retains the inertial and Coriolis coupling characteristic of multi-joint dynamics while omitting the actuator redundancy, muscle nonlinearity, sensorimotor delay and signal-dependent noise of biological limbs, and show that the network, trained exclusively on inverse dynamics data, can accurately predict the system’s forward evolution faster than in real time. We further show that the energy parameterization gives a lower median torque error than direct torque regression by an otherwise identical network, and that the inverse and forward computations transmit input noise as their analytical counterparts do. This work provides a first-principles basis for unifying forward and inverse internal models within a single computational architecture, and in doing so removes one formal argument for treating them as separate. Whether the nervous system exploits such a shared representation is an empirical question that these results do not settle; they do, however, sharpen it, because a shared acceleration-energy representation predicts that competence acquired in one direction should transfer to the other, whereas independently parameterized models predict no such transfer. We test this prediction in simulation and show that adaptation of the inverse dynamics to a changed pendulum carries over to forward simulation without forward-specific training.

**Fig. 1.**
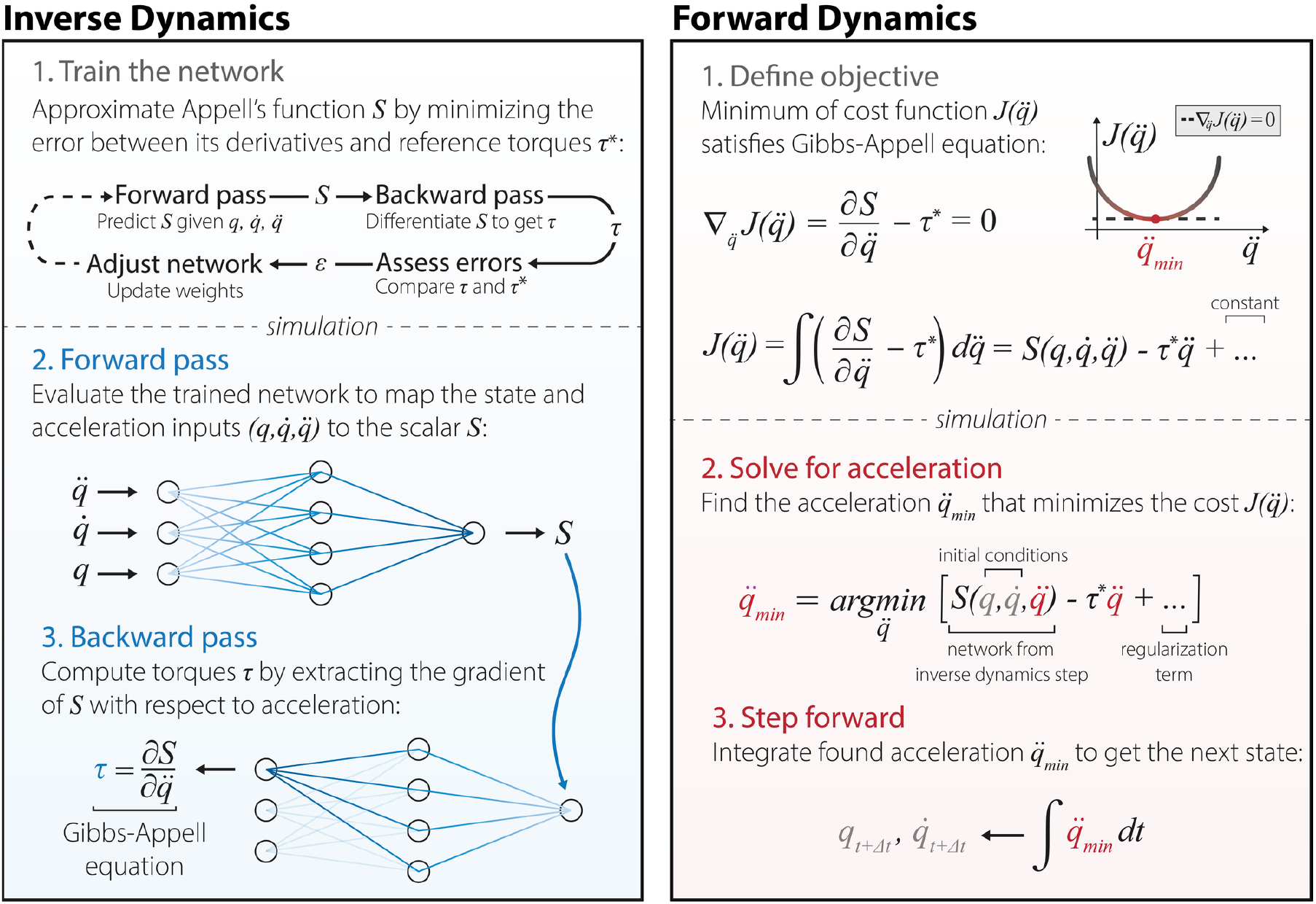
A single neural network trained to predict the Appell acceleration energy 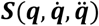 is shared between the inverse and forward dynamics pipelines. Left, a network learns the Appell acceleration energy, 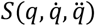, by minimizing the error between its analytical derivative and reference torques, *τ*\*, from analytical equations (the inertial torques of independently sampled states). In the simulation phase, the forward pass evaluates the network to map kinematic states 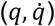 and accelerations 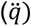 to the scalar *S*, while the backward pass computes torques (*τ*) for each joint by extracting the derivative of *S* with respect to acceleration 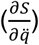 via the Gibbs–Appell equation. Right, optimization-based forward dynamics solver. A cost function, 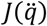, is formulated such that, prior to regularization, its unconstrained minimum analytically satisfies the Gibbs–Appell equation; in this panel *τ*\* denotes the prescribed gravitational forces, −**G**(**q**), held fixed during acceleration minimization step. To solve for acceleration, we isolate 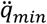 that minimizes cost 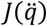, using the learned energy landscape from the inverse step alongside a temporal regularization term, before integrating the resultant acceleration to advance the state variables forward in time 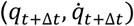.

## Methods

### Analytical formulation of system dynamics

Analytical dynamics of the system were formulated using Gauss’s principle of least constraint^48^. According to this principle, a constrained mechanical system evolves so that its actual accelerations deviate as little as possible from the unconstrained accelerations produced by applied active forces. Mathematically, this is achieved by minimizing the mass-weighted constraint function *Z* over the space of admissible accelerations:

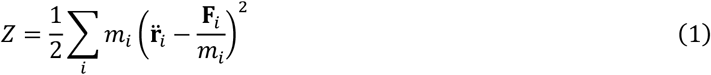

where *m*_*i*_ represents the mass of the *i*-th object, 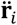 its actual acceleration, and **F**_*i*_ the applied active force. Expanding *Z* separates the function into three components:

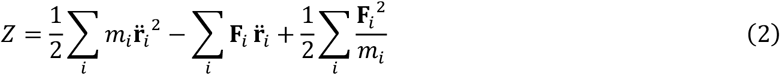

The first term, which is quadratic in acceleration, constitutes the Appell acceleration energy *S*^49,50^. The second term is linear in acceleration, representing the work-like contribution of active forces. The third term is independent of acceleration and is determined exclusively by the system’s masses and the applied forces (e.g., gravity). Because these applied forces are determined by the system’s current position and velocity rather than its instantaneous acceleration, this third term acts as a constant during the minimization process.

To find the minimum of the constraint function over the space of admissible accelerations, *Z* must be minimized with respect to the system’s independent generalized accelerations 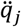, where *j* is the index of the degrees of freedom. This requires setting the partial derivative of *Z* with respect to each 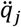 to zero:

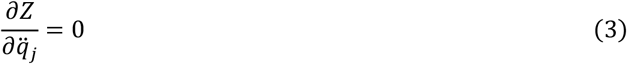

Because the third term in the expansion of *Z* is a constant with respect to 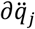, its derivative vanishes. However, minimizing *Z* is not equivalent to minimizing *S* alone because the active-force term remains. The stationarity condition is therefore:

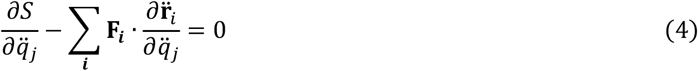

To evaluate the remaining sum, we apply the fundamental kinematic identity that relates Cartesian accelerations to generalized coordinates (a.k.a. ‘cancellation of dots’): 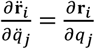. Substituting this identity into the minimization condition gives:

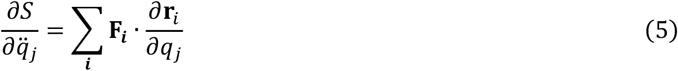

The right side of this expression constitutes the generalized active force *Q*_*j*_. Substituting this yields the final Gibbs–Appell equations of motion^51^:

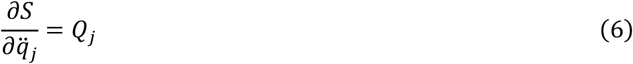

### Derivation of the double pendulum dynamics

We applied analytical derivations from above to model the dynamics of a double pendulum, representing it as two point masses, *m*_1_ and *m*_2_, within an inertial Cartesian frame. The physical connections between these masses (specifically, the massless rigid pendulum links of lengths *l*_1_ and *l*_2_) act as holonomic constraints that restrict the system’s motion. If we were to track the masses using Cartesian coordinates, we would have to solve a complex system of differential-algebraic equations to continuously enforce these fixed link lengths at every time step. Instead, we parameterized the system using a minimal set of independent generalized coordinates, **q** = [*θ*_1_, *θ*_2_]^⊤^, corresponding to the joint angles. Here, *θ*_1_ was the absolute first-link angle and *θ*_2_ the relative second-joint angle, so the absolute second-link angle was *θ*_1_ + *θ*_2_. Because these angles entirely define the system’s configuration, a forward kinematic mapping from **q** to the Cartesian position vectors **r**_*i*_ intrinsically satisfies the geometric constraints.

Next, taking the second time derivative of these position vectors symbolically in SymPy (v.1.14.0) yielded the Cartesian accelerations 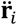 as explicit functions of the system’s joint angles, velocities, and accelerations 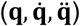. The Appell acceleration energy *S* was then constructed as half the mass-weighted sum of the squared Cartesian accelerations (nominal simulations used *m*_1_ = *m*_2_ = 1 kg, *l*_1_ = *l*_2_ = 1 m, *g* = 9.81 m/s^2^):

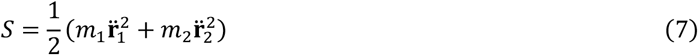

Because the system evolved solely under the influence of gravity, the active generalized forces driving the motion corresponded exclusively to the gravitational joint torques, τ ≡ − **G**(**q**). As dictated by the Gibbs–Appell formulation^52^, the physical system balances these active forces against the partial derivatives of the acceleration energy with respect to the generalized accelerations. Equating these terms yields the final equations of motion:

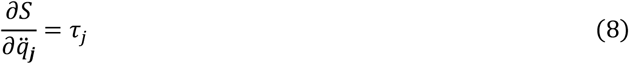

Numerically integrating these ordinary differential equations generated the reference trajectories used to evaluate the trained networks (see *Generation of ground-truth trajectories via forward dynamics*). Equation 8 describes gravity-driven motion; the training labels are defined from its left-hand side alone, evaluated at independently sampled states (see *Synthetic data generation and standardization*).

### Synthetic data generation and standardization

To generate the training dataset, we sampled kinematic states from uniform distributions, with joint angles *q*_*j*_ ∈ [−*π*/2, *π*/2] rad, velocities 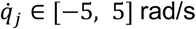, and accelerations 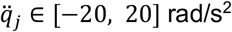, for the links *j* ∈ {1,2}. We chose these bounds to cover most of the angles, velocities and accelerations that the pendulum reaches under gravity (recognizing that independent, coordinate-wise uniform sampling defines an axis-aligned region rather than a rotationally invariant distribution in the six-dimensional state space).

The accelerations were sampled separately from the angles and velocities. When the pendulum swings freely under gravity, its angles and velocities already determine its acceleration, so data recorded along such swings would never show the network two different accelerations for the same angles and velocities. The forward solver, however, holds the angles and velocities fixed and compares many possible accelerations, and the network can only be trusted on such comparisons if it has seen similar combinations during training. For each sampled state vector 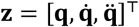, we computed the Appell acceleration energy *S* and the torques τ as the acceleration-gradient of *S* at the sampled state, 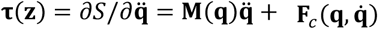, where **M**(**q**) is the configuration-dependent mass matrix and 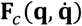 contains the velocity-dependent (Coriolis and centripetal) inertial terms (see *Supplementary Information, Forward dynamics equations*). Because 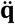 is sampled independently of 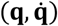, the sampled states do not lie on gravity-driven trajectories and ***τ*** (**z**) is in general different from −**G**(**q**); as such, the labels are the generalized inertial forces associated with the sampled acceleration, and gravity does not enter them. Equation 8 is therefore recovered on physical trajectories, where 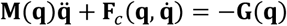.

We next standardized the input states **z** and the acceleration energy *S* to zero mean and unit variance, denoting these normalized, dimensionless variables as **z**\* and *S*\*. The target torques ***τ*** remained unnormalized. Since subsequent computational differentiation would evaluate the gradient of *S*\* with respect to the dimensionless accelerations 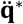, we computed and stored a chain-rule scaling vector, 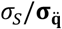. Here, the scalar *σ*_*S*_ and the two-element vector 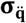 denote the sample standard deviations of the energy and the accelerations across the dataset, respectively, and provide the transformation required to map the dimensionless partial derivatives back to the physical domain, ensuring that 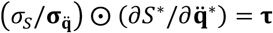, where ⊙ denotes the element-wise product.

### Neural network architecture

We designed a feedforward artificial neural network to map the normalized state vector **z**\* ∈ ℝ^6^ to the dimensionless Appell acceleration energy *S*\* ∈ ℝ. The finalized architecture, determined via the hyperparameter optimization (see *Hyperparameter optimization and architectural search*), comprised a multilayer perceptron with four hidden layers containing 225, 61, 91, and 99 computational nodes, respectively. We applied Gaussian Error Linear Unit (GELU) activation functions to each hidden layer^53^ and terminated the network with a single linear node to output the scalar acceleration energy. Batch normalization was not applied within the layers; instead, the six input coordinates were standardized using fixed dataset-level statistics before training.

### Physics-informed loss formulation

To train the network, we formulated a physics-informed loss function based on the Gibbs–Appell equations. Rather than computing a loss on the scalar acceleration energy output directly, we evaluated the network’s ability to predict joint torques. We chose this gradient-based supervision for two reasons. Theoretically, minimizing the error on a scalar function does not guarantee that a neural network will accurately capture the function’s gradients, which are required to represent the actual equations of motion. Practically, deriving the analytical Appell acceleration energy is cumbersome for complex, high-DOF systems, whereas joint torques are standard physical quantities that can be obtained readily via sensors or inverse dynamics solvers. Notably, the relevant constraint is not global recovery of the scalar acceleration-energy surface, but accuracy of the scaled input-gradient that enters the Gibbs–Appell equations. Our loss therefore supervises this derivative quantity directly, consistent with Sobolev training, where derivative information is incorporated into the learning objective^54^.

During each forward pass, we computed the numerical partial derivatives of the network’s output *S*_*net*_* with respect to the normalized input accelerations 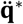 using automatic differentiation. We multiplied these dimensionless gradients by the precomputed scaling vector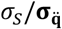 to predict torques, ***τ***_pred_. The torques stayed in physical units because they enter training only as loss targets, never as network inputs, and the chain-rule scaling already returns the predicted gradient to Nm. We defined the optimization objective as the mean squared error between ***τ***_pred_ and the ground-truth joint torques ***τ*** across the batch size *B* = 64:

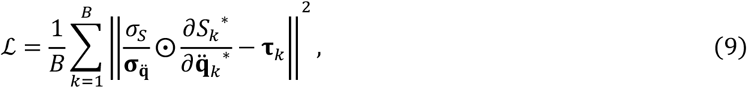

where *k* is the index variable for the individual data samples within a given mini-batch (a set of samples processed together).

### Hyperparameter optimization and architectural search

To identify the optimal network architecture and training hyperparameters, we conducted a systematic hyperparameter search in Optuna (v.4.8.0)^55^. We evaluated candidate configurations on an independent dataset of 10,000 kinematic samples (chosen to densely span the underlying kinematic phase space), partitioned into an 80% training set and a 20% validation set, applying the previously described standardization and chain-rule scaling transformations. The architectural search space included a discrete range of 1 to 10 hidden layers (with 1 to 256 nodes per layer) and a categorical selection of activation functions (GELU, Tanh, Softplus, ELU and ReLU). Finally, the training learning rate was sampled from a continuous log-uniform distribution between 10^−8^ and 10^−1^.

The search space was sampled across 500 independent trials, each representing a distinct parameter set. We evaluated each set of parameters for 100 epochs, using the physics-informed loss ℒ as the minimization objective. (We report the square root of the validation loss of each trial as its torque RMSE.) To allocate computational resources toward converging models, we employed a median pruning algorithm that terminated trials whose performance was worse than the historical median validation loss after a ten-epoch warmup phase. Of the 500 initiated trials, the pruner terminated 319. We quantified the share of the variation in the search objective attributable to each hyperparameter across the 181 trials that ran to completion with the functional analysis of variance (fANOVA) importance evaluator of Optuna^55^.

The algorithm converged on an optimal architecture of 225, 61, 91, and 99 nodes for the first, second, third, and fourth hidden layers, respectively, alongside GELU activations and a learning rate of 1.13 × 10^−3^. This optimal configuration served as the final architecture for all subsequent experiments.

### Experimental setup and optimization

To evaluate the data requirements of the architecture, independent models of the same architecture were trained on nine datasets doubling in size from 1,000 to 256,000 total vectors **z**\*. Each dataset was partitioned using the established 80% training and 20% validation split. The network weights were optimized using the Adam algorithm^56^ with the optimal learning rate (see *Hyperparameter optimization and architectural search*). Each model was trained for 3,000 epochs. Training loss was recorded every epoch and validation loss every 100 epochs to evaluate convergence, and inference timing was measured to assess the computational viability of the trained networks. For each dataset size, downstream analyses used the checkpoint with the minimum periodic validation error, rather than the weights from the final training epoch. The forward solver, the baseline comparison, the noise analyses and the transfer test used the model trained on 16,000 samples (see *Results* for its selection).

### Generation of ground-truth trajectories via forward dynamics

The inverse dynamics predictions of the trained networks were evaluated using a fixed test dataset comprising 50 independent temporal trajectories generated through numerical integration. The partial derivatives of the Appell acceleration energy were set equal to the gravitational forces derived from the potential energy *V* = *m*_1_ · *g*· *y*_1_(*θ*_1_, *θ*_2_)+ *m*_2_ · *g*· *y*_2_(*θ*_1_, *θ*_2_), yielding:

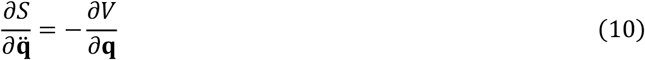

We algebraically inverted this system of equations to isolate the generalized accelerations 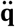 as explicit functions of the current state (*see Forward dynamics equations in the Supplementary information section*). To enable numerical integration, we reformulated the resulting dynamics as a system of continuous first-order ordinary differential equations. To establish the initial conditions, we sampled the joint angles uniformly from the interval [−*π*/3, *π*/3] rad and set the initial angular velocities to 0 rad/s, which ensured that the subsequent motion was driven exclusively by gravitational forces. Finally, we numerically integrated this system of equations over a five-second duration using a fourth-order Runge–Kutta scheme implemented in PyTorch with a fixed time step of 10^−4^ s (see ^57,58^ for similar solver configuration). Because the trajectories evolved freely under gravity, the evaluation states were not confined to the training sampling ranges. Across the 50 trajectories, 3.4% of states fell outside the sampled region on at least one coordinate (0% for both angles and for the velocity and acceleration of joint 1; 0.95% for the velocity of joint 2, which reached 7.4 rad/s; 2.7% for the acceleration of joint 2, which reached 36 rad/s^2^), affecting 9 of the 50 trajectories; no angle exceeded 1.3 rad, so no angle wrapped around the axis.

### Neural network-based inverse dynamics

We deployed the trained models to predict the inverse dynamics of the 50 unseen ground-truth trajectories. For each model, we normalized the kinematic state vectors of the test set using the mean and standard deviation parameters derived from its specific training distribution. The network mapped these standardized inputs to predict the dimensionless acceleration energy *S*\*. We evaluated the partial derivatives of this scalar output with respect to the normalized input accelerations via automatic differentiation. We then multiplied these dimensionless gradients by the model-specific scaling vector to recover the predicted physical joint torques:

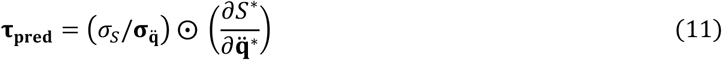

To quantify the predictive accuracy of each model, we computed the root-mean-square error (RMSE) between ***τ***_**pred**_ and the analytical ground-truth, evaluated for each trajectory and each joint across its 50,001 time steps.

### Ablation of the acceleration energy output

To test whether inverse-dynamics accuracy reflected the Gibbs–Appell energy parameterization rather than network capacity, we trained a direct torque regressor identical to ANNet except in its output layer. The network received the same standardized six-dimensional kinematic state and retained the four GELU hidden layers, but its output layer comprised two linear units that returned joint torques directly rather than the scalar acceleration energy, bypassing the automatic differentiation and gradient-rescaling steps of Eq. (11). We standardized the torque targets per joint using dataset-level statistics and de-standardized the network outputs before evaluating the loss, so that training minimized the mean squared error in physical units (the same quantity minimized by the physics-informed loss but requiring no input-gradient computation). All other settings were identical to those of the 16,000-sample ANNet used in the other analyses. We deployed this baseline on the same 50 unseen trajectories and computed the per-joint RMSE exactly as above.

### Sensitivity of the inverse model to input noise

We computed the torque error of the inverse model under Gaussian perturbation of its inputs to isolate its sensitivity to measurement noise. For each of the 50 test trajectories (50,001 samples per trajectory), we added zero-mean Gaussian noise independently to every sample of the two joint angles, velocities and accelerations, with a standard deviation of 1, 2.5, 5, 10 or 25% of the standard deviation of that signal over the trajectory. We rescaled one seeded realization per trajectory across levels, so that conditions differed only in noise amplitude. The network trained on 16,000 samples and the analytical inverse dynamics received identical noisy inputs, and we computed the RMSE of each against the noise-free analytical torques for every trajectory and joint; applying the noise to the angles, velocities or accelerations alone isolated the contribution of each input. For display, we selected the trajectory whose ANNet errors at 5% noise lay closest to the medians of both joints. We compared the two models at each noise level and joint with the tests described under Statistical analysis (*α* = 0.05, Benjamini–Hochberg correction across the ten comparisons) and with two-sided Wilcoxon signed-rank tests paired by trajectory. We compared the ANNet errors under each of the four noise conditions (angles, velocities, accelerations or all inputs) with the noise-free ANNet errors of the same trajectories by the same tests, with correction across the 40 comparisons of joint, condition and level.

First-order error propagation gave the law by which the torque error grows with input noise. Noise with standard deviation *pσ*_*j*_ on input *x*_*j*_, where *p* is the noise level and *σ*_*j*_ the standard deviation of that input over the trajectory, gives an exact model the error *RMSE*_*i*_(*p*) = *k*_*i*_*p* for joint *i*, with the gain 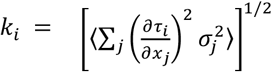 and the angle brackets denoting the average over time; a model whose noise-free error RMSE_*i*_(0) is independent of the noise follows *RMSE*_*i*_(*p*) = [*RMSE*_*i*_(0)^2^ + (*k*_*i*_*p*)^2^]^1/2^. We computed *k*_*i*_ for every trajectory by automatic differentiation of the analytical inverse dynamics on the noise-free inputs, and the same quantity from the Jacobian of ANNet, the second derivative of the learned energy. We quantified agreement as the relative deviation of the measured errors from each law and as the exponent of a power law fitted by least squares to the logarithms of error and noise level.

To measure the torque error that remains after the standard processing of measured kinematics, we low-pass filtered the noisy inputs before inverse dynamics. Each of the six noisy signals passed forward and backward through a second-order Butterworth filter (zero phase lag, fourth order overall; SciPy 1.16.3) at the sampling rate of 10 kHz, with mirror extension of the signal over three cutoff periods at both ends. The cutoff, defined as the –3 dB frequency of the two passes, was 1, 2, 5, 10, 20, 50, 100, 200, 500 or 1,000 Hz. ANNet and the analytical inverse dynamics received identical filtered inputs, and we computed the RMSE of each against the noise-free torques over the whole trajectory; filtering the noise-free inputs gave the error caused by the filter alone. We compared the ANNet errors with and without filtering by two-sided Wilcoxon signed-rank tests paired by trajectory, and the analytical errors with a first-order prediction, the root of the sum of the squared error caused by the filter alone and the squared product of the unfiltered error and the fraction of the white-noise standard deviation that the filter transmits.

### Forward dynamics via optimization

To solve the forward dynamics problem without requiring a distinct forward model, we repurposed the trained inverse dynamics network and formulated the derivation of joint accelerations as an unconstrained numerical optimization task. Following the Gibbs–Appell formulation, we established the fundamental equation of motion by equating the gradient of the acceleration energy with respect to the generalized accelerations to the generalized external forces. For the simplified double-pendulum system considered here, we set gravity as the only external generalized force; notably, this choice defines the test problem and is not a restriction of the network or optimization formulation. In systems with additional applied torques or external forces, these terms can be added to the generalized-force vector. Denoting gravitational contribution as τ ≡ −**G**(**q**), we obtained the governing equation of zero balance:

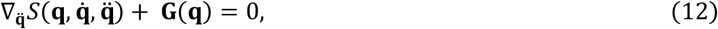

where 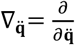. To resolve this equation algorithmically, we sought a scalar objective function 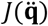 such that its unconstrained minimization (characterized by the stationary condition 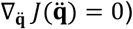 mathematically satisfies this balance of torques. We constructed 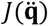 by taking the integral of the equation of zero balance with respect to the acceleration vector 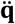:

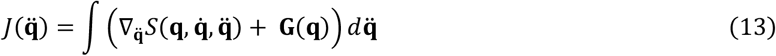

Because the inertial term is the gradient of the acceleration energy with respect to 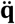, integrating it over acceleration space recovers *S* 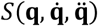, up to a term that may depend on **q** and 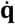 but not on 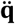. The gravitational vector **G**(**q**) is independent of acceleration, so its integral with respect to 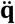 is the inner product 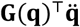. Combining these terms gives the objective function of form:

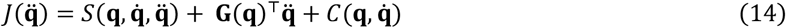

The aggregated integration constant 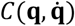 *does not* depend on the optimization variable 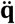. Defining *J*_0_ as the acceleration-dependent part of the preceding expression gives:

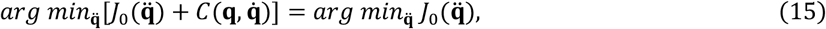

so, omitting 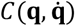 during the computation of the acceleration that minimizes the cost 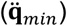 does not change the minimizing acceleration. As such, while retained for analytical correctness, term 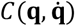 was omitted during the computational implementation.

To formulate the final objective function for simulation, we replaced the analytical *S* with the network-predicted acceleration energy *S*_*net*_. Because the network outputted a dimensionless, standardized scalar *S*_*net*_*, we converted the network output to acceleration energy units by applying the inverse Z-score transformation, *S*_*net*_ = *S*_*net*_\**σ*_*S*_ + *μ*_*S*_, using the precomputed dataset statistics. This denormalization ensured dimensional consistency across the objective function, to which we added a temporal regularization term:

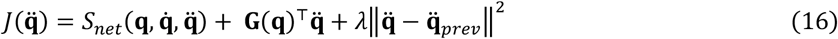

where 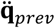 denotes the acceleration from the preceding time step, and *λ* = 0.1 is the regularization coefficient. Unlike the integration constant 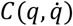, which was dropped because it does not depend on 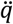, the regularization term depends on 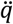 and can shift the minimizer away from the solution of the equation of zero balance.

We minimized the objective function at each timestep using a two-stage solver, with optimal parameters determined via grid search (see *Performance optimization of the optimization-based forward dynamics solver* section). In the first stage, a coarse global search evaluated the cost function across 16 candidate acceleration vectors. This candidate pool comprised the zero vector, the acceleration from the previous timestep, and 14 vectors randomly sampled from a uniform distribution bounded by ±15 rad/s^2^. In the second stage, the candidate yielding the minimum cost was used to initialize a gradient-based local refinement. We executed 10 optimization steps using the Adam algorithm (learning rate = 0.1) to converge on the final minimizer.

### Performance optimization of the optimization-based forward dynamics solver

The accuracy and efficiency of this two-stage solver depend on the scale of the candidate pool and the depth of the local gradient refinement. To systematically identify the optimal solver configuration, we performed a grid search across 336 distinct parameter sets. Our search evaluated three parameters, namely the number of sampled acceleration vectors (*N*), the uniform acceleration sampling bound (*B*), and the number of Adam refinement steps (*A*) (Table 1). We evaluated each configuration over 50 trajectories with 200 closed-loop state updates per trajectory, yielding 10,000 updates per configuration (see *Closed-loop simulation* for the initial-state ranges, the state update and the reference integration).

**Table 1.**
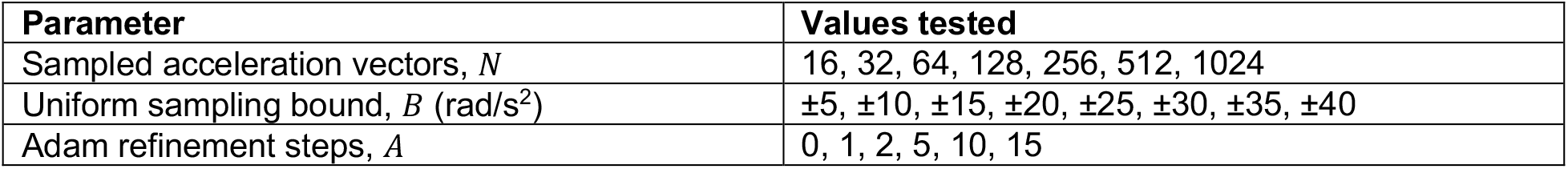
Parameter search space for the forward-dynamics solver.

Solver performance was assessed by joint angular accuracy and computational latency. The primary accuracy metric was the mean absolute error (MAE) across both joints, calculated per state update as:

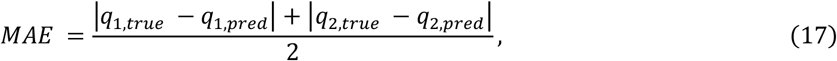

where *q*_1,*true*_ and *q*_2,*true*_ are the ground-truth angles for joint 1 and joint 2, and *q*_1,*pred*_ and *q*_2,*pred*_ are the corresponding predicted angles. This value was averaged across the 10,000 updates.

To assess computational speed, we recorded the full state-update time rather than isolated neural-network inference time. This metric accounts for optimizer execution, candidate evaluation, network forward passes, Adam refinement, numerical integration, tensor operations, and loop overheads. Given a simulation timestep of 2 ms, a configuration was defined as real-time capable if its mean update time was ≤2.0 ms.

### Closed-loop simulation

The predictive accuracy of the optimization-based forward dynamics solver was evaluated by conducting closed-loop forward simulations of 5 s from 50 random initial states, with the joint angles drawn uniformly from –0.5 to 0.5 rad and the angular velocities from –0.5 to 0.5 rad/s. For each initial state, the system was integrated forward in time using a semi-implicit Euler integration scheme (Δ*t* = 2 ms), chosen to maintain the numerical stability of the first-order update. At each temporal increment, the solver assessed the current state variables 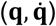 and computed the resultant acceleration vector, 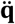. To stabilize the temporal regularization term at the onset of each simulation, a five-iteration stabilization loop was applied at *t* = 0 s, with the optimizer’s output recursively fed back to establish the boundary condition for 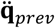.

Integration drift was quantified by generating corresponding ground-truth trajectories for identical initial states using an explicit fourth-order Runge–Kutta integration scheme evaluated on the same 2,500-point time grid as the closed-loop simulation (Δ*t* = 2 ms) (see ^59^ for similar solver configuration). All states of these reference trajectories remained within the training sampling ranges. The continuous state evolution was recorded, and the RMSE was computed between the simulated and ground-truth angles, velocities, and accelerations across the temporal horizon for both joints.

### Isolating solver and integrator error

To separate the error contributed by the learned energy from that contributed by the optimizer and the integrator, we repeated the closed-loop simulation with the analytical Appell acceleration energy 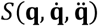 substituted for *S*_*net*_ in the objective function from Eq. 16. The substitution replaced the energy and its gradient together, so the refinement steps were driven by the exact analytical 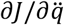 rather than by automatic differentiation through the network; the residual error in this condition therefore reflects the candidate search and refinement, the temporal regularizer and the integration scheme. All other components of the solver were unchanged. We integrated the resulting trajectories with a semi-implicit Euler scheme (Δ*t* = 2 ms) for 5 s from the same 50 initial states used in the learned-energy run, compared them with the corresponding fourth-order Runge–Kutta reference trajectories, and computed the RMSE of joint angles, velocities and accelerations per trajectory and joint as described above.

### Sensitivity of the forward solver to noise in the gravitational torques

We computed the error of the closed-loop forward simulation under Gaussian perturbation of its input, the gravitational torque, to isolate the sensitivity of the forward solver to noise in the generalized forces. At every solver step we added zero-mean Gaussian noise to the gravitational torque of each joint, independently across joints and steps, with a standard deviation of 1, 2.5, 5 or 10% of the standard deviation of that torque over the noise-free reference trajectory; the initial angles and velocities remained exact. The simulation protocol was otherwise that of the 50 noise-free closed-loop simulations (see *Closed-loop simulation*), with one seeded noise realization per trajectory rescaled across levels and identical candidate accelerations in every condition, and we recomputed the noise-free simulation with the same code. The solver with the analytical Appell energy received the same noise realizations, and we computed the RMSE of angle, velocity and acceleration of each solver against the noise-free fourth-order Runge–Kutta reference for every trajectory and joint. We compared the two solvers without noise and at each noise level (30 comparisons), and each noise level with the noise-free simulation (24 comparisons per solver), with the tests described under *Statistical analysis* and with two-sided Wilcoxon signed-rank tests paired by trajectory (*α* = 0.05, Benjamini–Hochberg correction within each family).

### Transfer of inverse-dynamics learning to forward dynamics

To isolate the learning that reaches the forward dynamics through the shared acceleration energy, we adapted the network to perturbed pendulums with the physics-informed loss ℒ alone. We changed the point masses from 1 and 1 kg to 1.2 and 0.8 kg, the link lengths from 1 and 1 m to 0.9 and 1.1 m, or both, and drew 16,000 states of each perturbed pendulum from the original sampling ranges (80% for training), with its analytical torques as targets. The network trained on 16,000 samples of the original pendulum (the pretrained network) trained on these data for 300 epochs with the Adam optimizer, learning rate, batch size and standardization constants of the original training, so that the torque error was the only training signal; as a control, the pretrained network received the same training on fresh samples of the original pendulum (the control network). We stored the weights before adaptation and after 1, 3, 10, 30, 100 and 300 epochs and defined the network of epoch 300 as the adapted network, whereas the pretrained network was the checkpoint with the lowest validation loss of its 3,000 training epochs.

Closed-loop forward simulations with every stored network gave the forward accuracy at each stage of this training. The forward solver, in the configuration selected above and supplied with the gravitational torques of the simulated pendulum in every condition, simulated each pendulum for 5 s from the 50 initial states with each stored network and with the analytical acceleration energy of that pendulum. We refer to the solver with the adapted, control or pretrained network as the adapted, control or pretrained solver, to the solver with the analytical energy as the analytical solver, and to the pretrained solver with the gravitational torques of the original pendulum as the model without any update. We computed the RMSE of angle, velocity and acceleration against fourth-order Runge–Kutta reference trajectories of the simulated pendulum, the same RMSE of the model without any update against the reference of each perturbed pendulum, and the RMSE of the network torques on the reference trajectories against the analytical inverse dynamics.

We tested the change of error with adaptation and the distance of the adapted solver from four references. We compared the errors at each stored epoch with those before adaptation (108 forward and 36 torque comparisons for the perturbed pendulums, 36 and 12 for the original pendulum). We compared the forward errors after 300 epochs with those of the analytical solver (24 comparisons, 18 for the perturbed pendulums and 6 for the control) and, with 18 comparisons each, with those of the control solver, of the pretrained solver on the original pendulum and of the model without any update. Each comparison used the unpaired tests described under *Statistical analysis* and a two-sided Wilcoxon signed-rank test paired by trajectory (*α* = 0.05, Benjamini–Hochberg correction within each family).

### Statistical analysis

All statistical analyses were performed using SciPy^60^ (v.1.16.3) and statsmodels^61^ (v.0.14.6), while neural network computations and automatic differentiation were executed via PyTorch^62^ (v.2.11.0). All computations ran in Python (v.3.13.11). No trajectories were excluded. We evaluated the assumption of normality for all empirical distributions using the Shapiro–Wilk test (*α* = 0.05). Because the distributions of latencies and of torque RMSE across dataset sizes deviated from normality, we used non-parametric tests for these comparisons. One-sided Mann–Whitney U tests were used for directional latency comparisons when the prespecified hypothesis was that the optimization-based forward dynamics solver required longer execution time, measured as full-update time, than the traditional, numerical solver. In the baseline, analytical-energy, noise and transfer analyses, two groups were compared by one-way ANOVA when both passed the Shapiro–Wilk test and by the Kruskal–Wallis test otherwise, with Benjamini–Hochberg correction within each family of comparisons. We also compared the errors paired by trajectory with two-sided Wilcoxon signed-rank tests and report Cliff’s *δ* as the effect size in the baseline and analytical-energy comparisons. Adjusted *p* values are written *p*\*.

To map the statistical effect of training dataset volume on predictive accuracy, we performed pairwise comparisons of the RMSE distributions across dataset sizes using two-sided Mann–Whitney U tests. These pairwise comparisons were corrected for multiple comparisons using the Benjamini–Hochberg false discovery rate (FDR) procedure. We described the dependence of the median RMSE of each joint on dataset size with a power law plus a constant offset, fitted by non-linear least squares with non-negative coefficients.

To quantify computational efficiency and eliminate hardware biases, models were evaluated in a randomized, interleaved order with hardware operations synchronized immediately before and after execution. Timing summaries distinguish three distinct quantities. Network-forward time isolates the duration required for neural-network forward evaluations. Solver-call time encompasses candidate sampling, candidate scoring, and Adam refinement for a single optimizer call. Full-update time is the total wall-clock rollout time divided by the number of simulated state updates. Because the full-update metric includes the complete simulation loop overhead, it serves as the most conservative timing measure and was exclusively used to evaluate real-time performance. Differences in inverse-model inference latency across dataset sizes were evaluated using a Kruskal–Wallis H-test.

### Hardware specifications

All computational experiments, including dataset generation, network training, and dynamics simulations, were performed locally on an Apple M4 system-on-a-chip equipped with 32 GB of unified memory, running macOS Tahoe (v.26.3.1) for the hyperparameter search, the training across dataset sizes, the solver grid search, the closed-loop simulations and the timing measurements, and macOS (v.27.0) for the baseline, analytical-energy, noise and transfer analyses. Computations were executed exclusively on the CPU cores, without utilizing the integrated Metal Performance Shaders (MPS) for hardware acceleration.

## Results

### Hyperparameter search optimizes torque recovery from learned Appell energy

We selected the network architecture using inverse-dynamics accuracy as the performance criterion. Each candidate network received the standardized six-dimensional input vector **z**\* and returned a scalar estimate of Appell acceleration energy, *S*\*. To evaluate each candidate, we differentiated this scalar output with respect to the acceleration inputs and rescaled the derivative to obtain predicted joint torques. We therefore ranked the 500 hyperparameter trials by torque RMSE, directly testing the quantity required for inverse dynamics. Global sensitivity analysis showed that performance was most strongly affected by the width of the first hidden layer and the learning rate, which accounted for 51.4% and 25.5% of the variation in the search objective, respectively. Network depth accounted for 18.3%, whereas activation function accounted for 4.8%, indicating smaller but measurable effects (Fig. 2h).

**Fig. 2.**
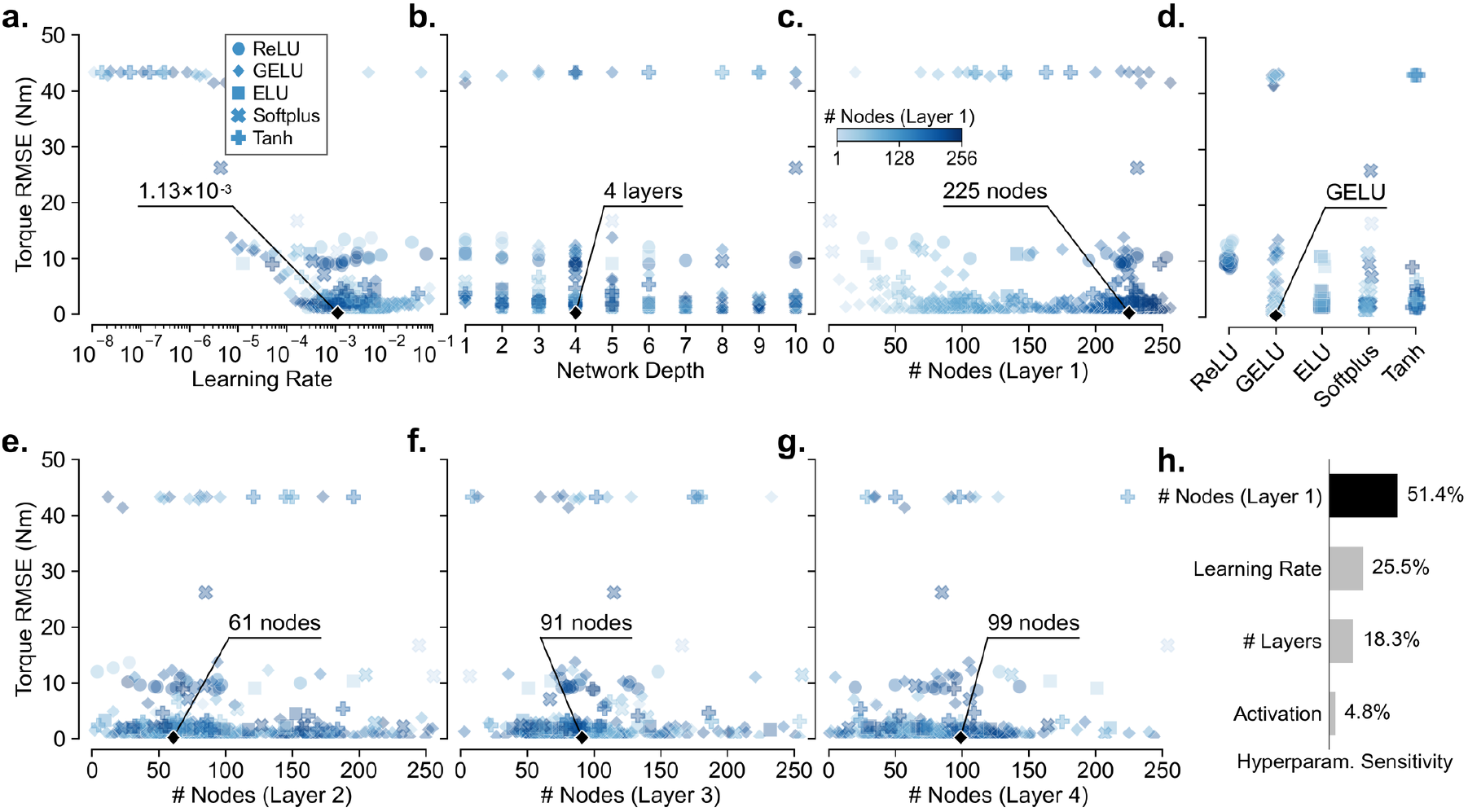
Systematic hyperparameter optimization. **a–g**, Torque prediction error for individual hyperparameter-optimization trials, reported as root-mean-square error (RMSE; Nm), plotted against learning rate (**a**), network depth (**b**), number of nodes in hidden layer 1 (**c**), activation function (**d**), and number of nodes in hidden layers 2–4 (**e–g**). Each symbol represents one evaluated model configuration (*n* = 500 trials, with the panels for hidden layers 2–4 restricted to the 481, 404 and 345 trials whose networks had that layer); symbol shape denotes the activation function, as indicated in **a**, and symbol color denotes the number of nodes in hidden layer 1, from light blue (1 node) to dark blue (256 nodes), as indicated in **c**. Symbols are drawn semi-transparent, so overlapping trials appear darker. Black diamonds mark the best-performing configuration, with learning rate = 1.13 × 10^−3^, network depth = 4 layers, GELU activation, and 225, 61, 91 and 99 nodes in hidden layers 1–4, respectively. **h**, Relative hyperparameter sensitivity, showing the proportional contribution of each parameter to variation in optimization performance across the *n* = 181 trials that ran to completion. The black bar marks the number of nodes in hidden layer 1, the hyperparameter encoded by the symbol color in **a–g**.

The search revealed limits on the hyperparameter values that produced accurate models. Networks trained with learning rates below 10^−5^ did not converge and remained at high torque errors near 43 Nm (Fig. 2a).

Lower errors required learning rates near 10^−3^. Accuracy also depended strongly on the width of the first hidden layer. The best-performing trials used first-layer widths close to the upper end of the search range, 225 nodes (Fig. 2c). In contrast, later hidden layers were less sensitive to width and reached low errors with fewer units (Fig. 2e–g). The lowest torque RMSE was obtained with a four-hidden-layer network, indicating that this depth performed best among the tested architectures (Fig. 2b).

Although the activation function had a smaller overall effect than learning rate or first-layer width, it still influenced the minimum error achieved by each model (Fig. 2d). Networks with GELU activations achieved lower torque RMSE than networks using ReLU, ELU, Softplus, or Tanh. This result is consistent with the derivative-based training objective, because torque predictions depend on the derivative of the learned scalar *S*\* with respect to acceleration. As such, across the full search, the best-performing configuration used a learning rate of 1.13 × 10^−3^, four hidden layers with 225, 61, 91, and 99 units, and GELU activations (Fig. 2a–g, black diamonds). We used this architecture for all subsequent analyses. A condensed parallel-coordinate representation of the full hyperparameter search is provided in Supplementary Fig. 1.

### Inverse-dynamics accuracy reaches a low error regime at intermediate dataset size

To determine how much training data the network requires, we trained the selected architecture on nine datasets of 1,000 to 256,000 samples and scored each model on the 50 test trajectories. Increasing the training set generally reduced inverse-dynamics error, but the improvement was not monotonic (Fig. 3). This non-monotonicity is expected under the fixed-architecture protocol used here. The architecture and learning rate were optimized once on an independent 10,000-sample architecture-search dataset and then held constant for all dataset-size experiments, so each point reflects data scaling within the same model class rather than a separately tuned model. Re-optimizing the architecture for every dataset size would conflate data efficiency with model selection, and the small reversals likely reflect finite held-out-set variation plus interactions between fixed model capacity, learning rate, training duration, and each training set. Across dataset sizes, validation RMSE declined rapidly during early training and then fluctuated within a lower-error regime rather than converging smoothly (Fig. 3a). These gains also came with a steep increase in computational cost, as training time rose from ~30 s for 1,000 samples to ~2.2 h for 256,000 samples (Fig. 3d, lower panel).

**Fig. 3.**
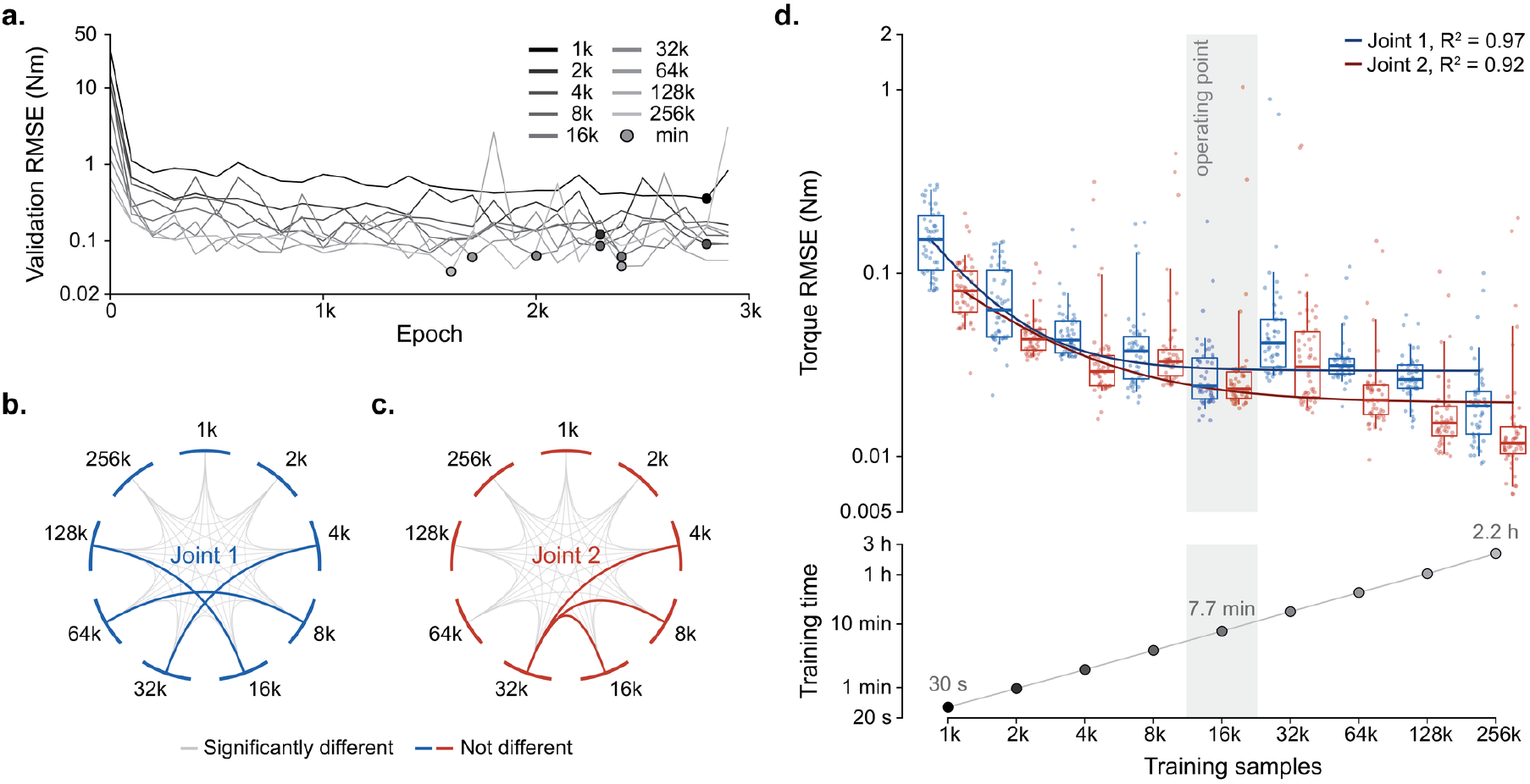
Dataset-size effects on inverse-dynamics accuracy and training cost. **a**, Validation RMSE during training for fixed-architecture networks trained for 3,000 epochs on datasets ranging from 1,000 to 256,000 samples (darker lines, smaller datasets). Markers indicate the validation-minimum checkpoint selected for downstream analyses. **b**,**c**, Pairwise comparisons of RMSE distributions (*n* = 50 test trajectories per dataset size) across dataset sizes for joint 1 (**b**) and joint 2 (**c**), computed using two-sided Mann–Whitney U tests with Benjamini–Hochberg correction. Grey chords connect dataset sizes whose distributions differed significantly and colored chords connect those that did not. **d**, Upper panel, torque RMSE distributions for joint 1 (blue) and joint 2 (red), evaluated on *n* = 50 independent 5-s test trajectories at each dataset size. Points show errors for individual trajectories, boxes indicate the IQR, centre lines indicate medians, and whiskers indicate the 5th–95th percentile range. Solid lines show power-law fits to the median RMSE values, with *R*^2^ values indicated. Lower panel, wall-clock training time for the same dataset sizes. The shaded band marks the 16,000-sample operating point used for subsequent analyses. All axes are logarithmic and span the full range of the plotted values.

Held-out performance showed a similar scaling pattern. The steepest gains occurred below 4,000 samples (3.5-fold and 2.8-fold reduction of median RMSE from 1,000 to 4,000 samples for joints 1 and 2); beyond 4,000 samples gains continued at a smaller and irregular rate, reaching 0.0242 and 0.0234 Nm at 16,000 samples, which captured 96% and 83% of the total reduction observed up to 256,000 samples for joints 1 and 2 (Fig. 3d, upper panel). Joint 2 error increased slightly between 4,000 and 8,000 samples, and both joints showed a transient increase in error at 32,000 samples before the downward trend resumed at larger dataset sizes. The lowest median errors occurred at 256,000 samples, reaching 0.0189 Nm for joint 1 and 0.0118 Nm for joint 2. Thus, larger datasets improved the best attainable accuracy, but much of the gain had already emerged by the intermediate dataset size regime. We therefore used the 16,000-sample model as the operating point for subsequent analyses because it captured most of the accuracy improvement while avoiding the substantially longer training times required at larger dataset sizes. To locate the largest errors of this model, we separated the test states inside the sampled ranges from those outside them. The largest trajectory-wise errors coincided with excursions beyond the sampled ranges. The mean absolute torque error was 0.022 and 0.020 Nm (joints 1 and 2) on the 96.6% of states inside the sampled region but 0.080 and 0.159 Nm on the states outside it, and the 41 trajectories that remained inside had maximum RMSEs of 0.040 and 0.033 Nm, whereas the 9 trajectories with excursions produced the outlying values in Fig. 3d, upper panel.

Consistent with the diminishing gains at larger datasets, the aggregate dependence of error on dataset size was well described by power-law fits to the median RMSE values, *RMSE*_*J*1_(*n*) = 21,782.3 · *n*^−1.75^ + 0.0293 (*R*^2^ = 0.974) for joint 1 and *RMSE*_*J*2_(*n*) = 108.958 · *n*^−1.089^ + 0.0195931 (*R*^2^ = 0.921) for joint 2, where *n* denotes the number of training samples. RMSE distributions were non-normal at all dataset sizes (Shapiro–Wilk, all *p* < 0.05), so group comparisons were performed using two-sided Mann–Whitney U tests with Benjamini–Hochberg correction shown as chord diagrams in Fig. 3b,c. An equivalent matrix representation of these FDR-corrected pairwise comparisons is provided in Supplementary Fig. 2. Relative to the 1,000-sample model, the 16,000-sample model significantly reduced error for both joints (joint 1, *p* = 8.48 × 10^−17^, *p*\* = 7.64 × 10^−16^; joint 2, *p* = 2.65 × 10^−14^, *p*\* = 2.01 × 10^−13^), supporting its use as an efficient operating point for downstream analyses.

To test whether this accuracy arose from the Appell-energy parameterization rather than from network capacity, we trained a network that differed from ANNet only in predicting the two joint torques directly from a linear output layer, holding architecture, data volume and training protocol constant. Under this identical budget the network mapping kinematics to torques directly converged to a higher minimum validation loss (9.5 × 10^−3^ versus 3.9 × 10^−3^ *Nm*^2^) and was less accurate on the held-out trajectories (Fig. 4). For joint 1, the median torque RMSE of the direct regressor was 0.0934 Nm compared with 0.0242 Nm for ANNet, a 3.9-fold increase (*H* = 66.40, *p* = 3.68 × 10^−16^, *p*\* = 7.37 × 10^−16^; Cliff’s *δ* = 0.95; paired Wilcoxon *p* = 4.91 × 10^−12^), and the network mapping to torques directly was less accurate in 49 of the 50 trajectories. For joint 2, the difference was smaller but remained significant (0.0288 versus 0.0234 Nm, a 23% increase; *H* = 10.86, *p*\* = 9.83 × 10^−4^; Cliff’s *δ* = 0.38; paired Wilcoxon *p* = 6.72 × 10^−4^; 39 of 50 trajectories). ANNet’s distributions contained a few high-error trajectories (up to 1.03 Nm for joint 2) that the direct regressor did not show, but these did not affect the median-based comparison. As such, with network capacity, data and optimization held constant, supervising the torques through the acceleration-gradient of a learned scalar energy yielded lower median torque error on the held-out trajectories than regressing the torques directly.

**Fig. 4.**
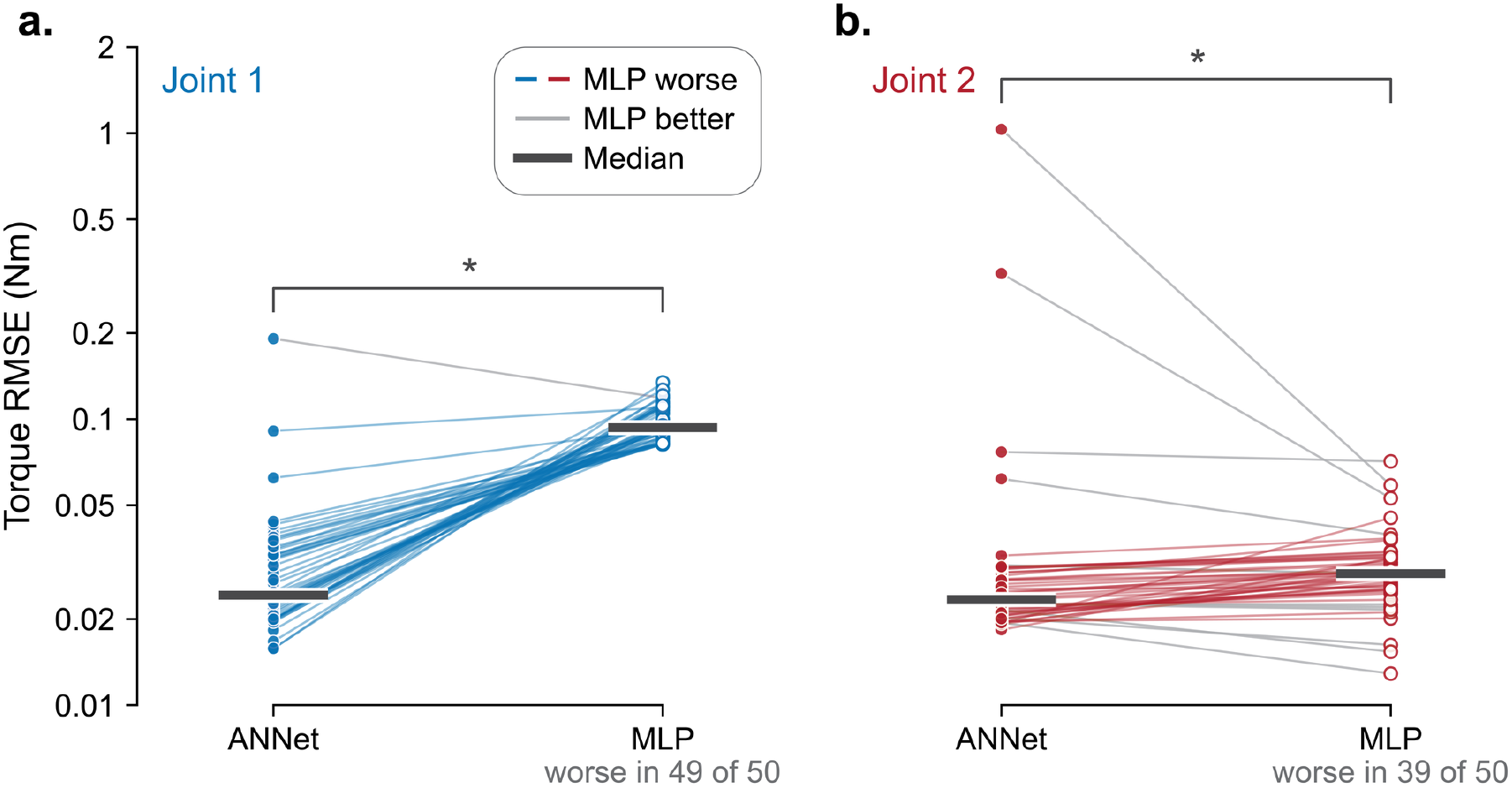
ANNet achieves a lower median torque error than an otherwise identical direct regressor. **a**,**b**, Torque RMSE of ANNet (filled circles) and of the direct regressor (open circles; labelled MLP, for multilayer perceptron) for joint 1 (**a**, blue) and joint 2 (**b**, red). The direct regressor matches ANNet in architecture and training protocol and differs only in its output layer, which returns the two joint torques directly. Points show errors for individual held-out 5-s test trajectories (*n* = 50 trajectories, each evaluated by both networks), and each line joins the two errors obtained on one trajectory. Lines are blue (**a**) or red (**b**) where the direct regressor had the higher error (MLP worse; the number of such trajectories is given beneath each panel) and grey where ANNet had the higher error (MLP better). Horizontal bars indicate medians, and the logarithmic axis is common to both panels. Asterisks mark differences between the two networks with *p*\* < 0.05, where *p*\* is the *p* value of a Kruskal–Wallis test, which treats the two sets of errors as independent samples, after Benjamini–Hochberg correction across the two joints (joint 1, *p*\* = 7.37 × 10^−16^; joint 2, *p*\* = 9.83 × 10^−4^). Two-sided Wilcoxon signed-rank tests, which account for the pairing by trajectory, also gave *p* < 0.05 for both joints (joint 1, *p* = 4.91 × 10^−12^; joint 2, *p* = 6.72 × 10^−4^; unadjusted).

Inference latency was effectively independent of dataset scale. Across 10,000 interleaved executions per model, median latency ranged from 73.29 *μ*s to 73.38 *μ*s, with IQRs of 1.59-1.71 *μ*s. Although latency distributions differed across dataset sizes (Kruskal–Wallis, *H* = 28.6616, *p* = 3.63 × 10^−4^), the total span of median latencies was only 0.083 *μ*s, indicating no practically meaningful deployment penalty from training on larger datasets.

### Forward-solver performance depends mainly on refinement depth and search bound

To identify the solver settings that recover accelerations accurately within the real-time limit, we ran the forward solver, built on the 16,000-sample network, over a grid of settings. Across 336 solver configurations, accuracy and latency formed a Pareto trade-off (Fig. 5). Configurations with little or no Adam refinement were fast but often inaccurate, whereas additional refinement reduced angular error at the cost of longer update times. Across the grid, median angular error decreased from 2.65 × 10^−2^ rad with no Adam steps to 8.37 × 10^−4^ rad with 10 Adam steps, while median update time increased from 0.22 ms to 1.73 ms. Increasing refinement to 15 steps produced only a small further reduction in error but raised the median update time above the 2.0-ms real-time limit.

**Fig. 5.**
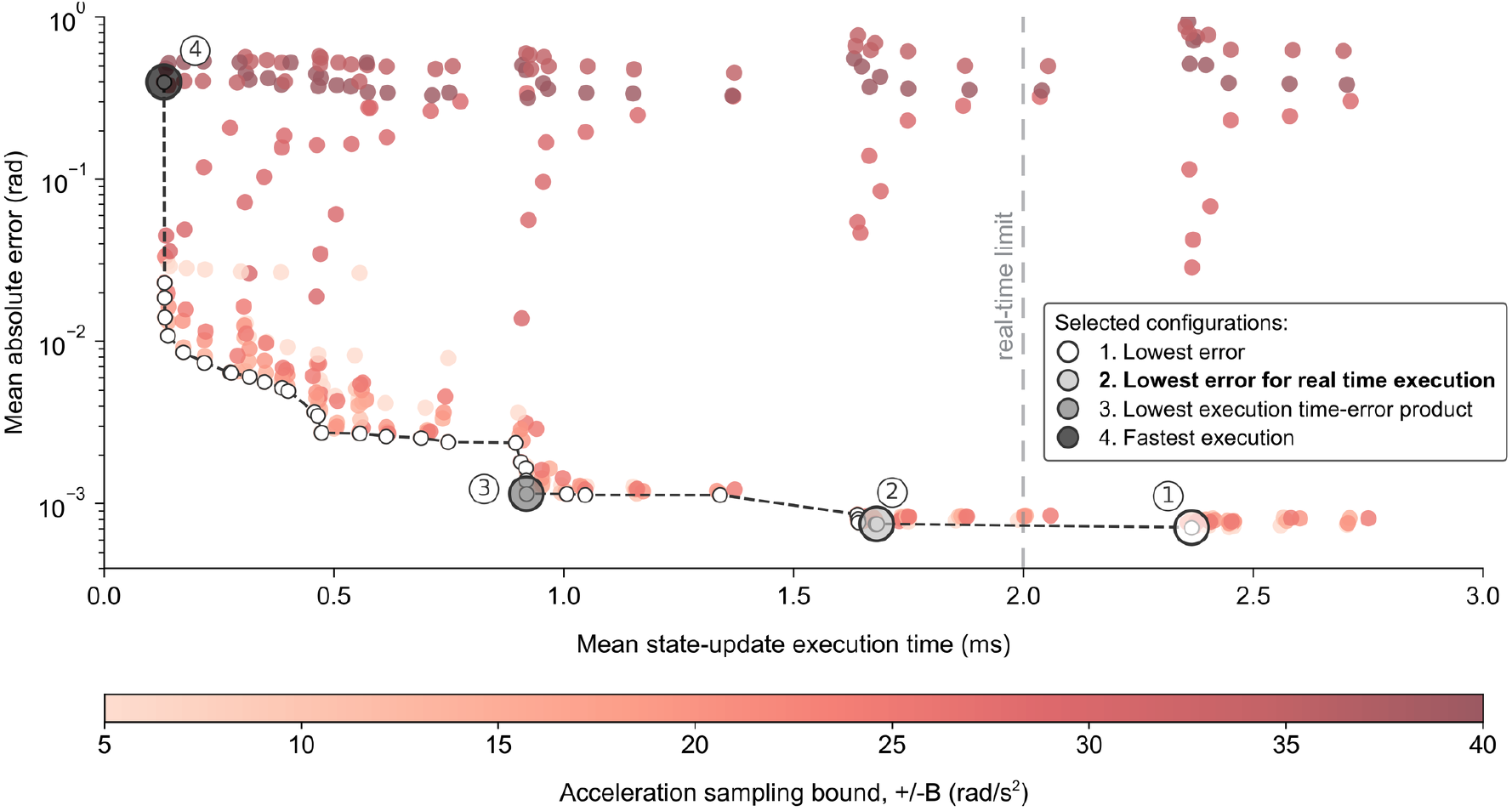
Accuracy-latency trade-offs of the optimization-based forward dynamics solver. Each point denotes one of 336 solver configurations, evaluated over *n* = 50 trajectories (10,000 closed-loop state updates). The y axis shows the mean absolute angular error, averaged across the two joints and plotted on a logarithmic scale; the x axis shows the mean state-update execution time. The grid varied the number of sampled acceleration vectors (*N*), the uniform acceleration bound (*B*, in ± rad/s^2^) and the number of Adam refinement steps (*A*); only *B* is encoded visually, from smaller bounds in light red to larger bounds in dark red. The black dashed curve joins the configurations on the empirical Pareto frontier (small white circles), and the grey dashed vertical line marks the 2.0-ms real-time limit. Numbered callouts and large semi-transparent circles identify the lowest-error configuration (1, white; *N* = 16, *B* = ±20, *A* = 15), the selected real-time operating point (2, light grey; *N* = 16, *B* = ±15, *A* = 10), the minimum execution-error product configuration (3, dark grey; *N* = 32, *B* = ±10, *A* = 5) and the fastest configuration evaluated (4, black; *N* = 16, *B* = ±35, *A* = 0).

The acceleration sampling bound also affected solver reliability. Moderate bounds produced the lowest-error operating points, whereas very broad bounds generated many high-error configurations, especially when local refinement was limited. The lowest-error configuration used 16 sampled acceleration vectors, a ±20 rad/s^2^ sampling bound, and 15 Adam steps, reaching a mean absolute angular error of 7.12 × 10^−4^ rad but requiring 2.37 ms per update. We therefore selected the best real-time-capable configuration, namely 16 sampled vectors, ±15 rad/s^2^ bound, and 10 Adam steps. This setting preserved nearly the same accuracy (7.50 × 10^−4^ rad) while reducing the mean update time to 1.68 ms. A faster balanced setting reduced update time to 0.92 ms with slightly higher error (1.15 × 10^−3^ rad), whereas the fastest configuration completed updates in 0.13 ms but had substantially larger error (0.397 rad).

### Learned Appell energy landscape enables accurate, real-time forward dynamic simulation

The selected optimization-based neural forward dynamics solver configuration (*N* = 16, *B* = ±15 rad/s^2^, *A* = 10) was tested in closed-loop forward simulation to determine whether the energy learned from inverse dynamics alone supports forward dynamics. The solver was evaluated on 50 reference trajectories, each lasting 5 s. At each time step, it estimated acceleration by first scoring 16 candidate acceleration vectors within the prescribed bound and then refining the best candidate with 10 Adam steps. The objective combined the learned Appell-energy term, the gravitational work term and a penalty for changes from the previous acceleration estimate.

The optimization-based solver closely followed the reference motion over the 5-s rollouts (Fig. 6a). Median angular RMSEs were 0.00737 rad for joint 1 and 0.0119 rad for joint 2. Median velocity RMSEs were 0.0309 and 0.0677 rad/s, and median acceleration RMSEs were 0.165 and 0.382 rad/s^2^, respectively. Errors accumulated gradually during the rollout, as expected for a closed-loop simulation that repeatedly feeds back its own acceleration estimates, but remained bounded over the full simulation horizon (Fig. 6b).

**Fig. 6.**
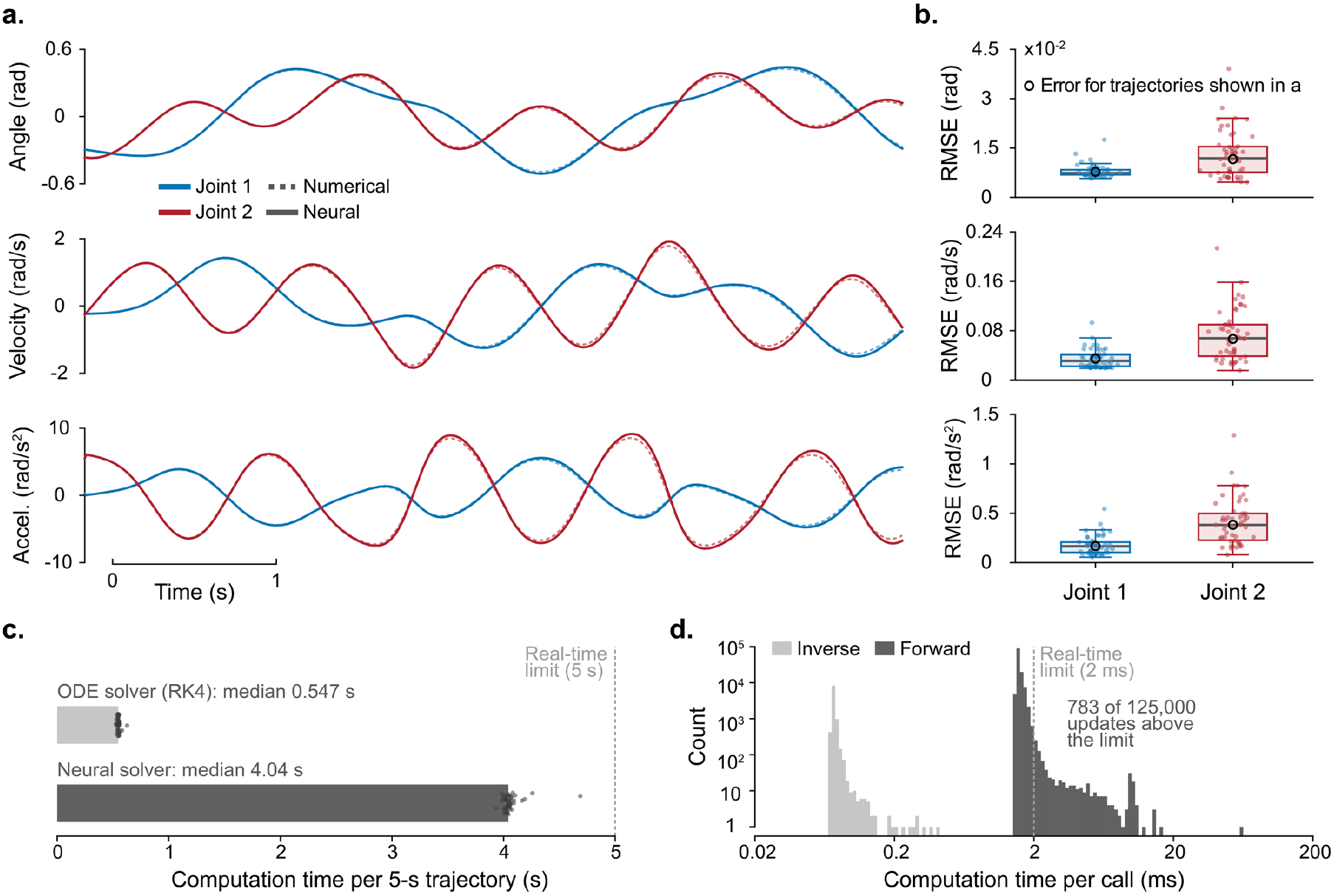
Learned acceleration energy landscape enables accurate and fast forward simulation. **a**, Representative closed-loop rollouts for the two-joint system, comparing the optimization-based neural forward dynamics solver (solid) with the numerical ODE solution (dashed) for angular position, velocity and acceleration. Blue and red denote joints 1 and 2, respectively. **b**, Trajectory-wise RMSE distributions for position, velocity and acceleration across *n* = 50 independent simulations, shown separately for each joint. Points denote individual trajectories; boxes span the interquartile range, centre lines denote medians and whiskers extend to 1.5 times the interquartile range. The open circle marks the trajectory shown in a. **c**, Computation time per 5 s trajectory for the numerical ODE solver and for the optimization-based neural forward dynamics solver; bars denote medians, points individual trajectories (*n* = 50 independent simulations per solver) and the dashed line the 5 s real-time limit. **d**, Distribution of per-query computation time for one inverse-model torque prediction (light grey, *n* = 10,000 queries) and one forward-solver update (dark grey, *n* = 125,000 updates); the dashed line denotes the 2-ms real-time limit, which only 783 of 125,000 updates exceeded; axes are logarithmic.

This accuracy came with a computational cost relative to direct numerical evaluation. For complete 5-s trajectories, the optimization-based solver required a median of 4.04 s, compared with 0.547 s for the numerical solver (Fig. 6c, one-sided Mann–Whitney *U* test, *U* = 2500, *p* = 3.53 × 10^−18^). At the individual update level, however, the solver remained within the 2.0-ms real-time budget. Median solver latency was 1.54 ms with an interquartile range of 0.0655 ms, and the mean update time over full rollouts was 1.62 ms (Fig. 6d). Thus, the learned Appell-energy landscape, while remaining slower than the traditional numerical integrator for complete trajectory generation, enabled accurate, real-time-capable forward simulation.

To determine how much of the closed-loop error originated in the learned energy rather than in the solver and the integrator, we repeated the simulation with the analytical Appell acceleration energy in the objective, holding the solver configuration, the initial states and the reference trajectories unchanged. Median RMSE in the run with the analytical energy term differed from the learned-energy run by at most 8% across joint angles, velocities and accelerations at both joints (ratios of medians 0.92–1.01; Cliff’s |*δ*| ≤ 0.07; Wilcoxon signed-rank, all *p* ≥ 0.41, *n* = 50 trajectories; Supplementary Fig. 3). Within the sampled state distribution, the closed-loop error of the forward solver was hence set by the finite candidate search and refinement schedule, the temporal regularizer and the first-order integrator rather than by the accuracy of the learned energy. The inverse and forward computations received exact inputs in all tests above, and the following sections examine them under noise and under a change of the pendulum.

### Input noise propagates through ANNet as through the analytical equations

To test whether the learned energy amplifies measurement noise beyond the amplification intrinsic to inverse dynamics, we compared ANNet with the analytical inverse dynamics on identical noisy inputs (Fig. 7a,b). We added Gaussian noise with a standard deviation of 1, 2.5, 5, 10 or 25% of the standard deviation of each input to the angles, velocities and accelerations of the 50 test trajectories and scored both models against the noise-free torques. The median torque RMSE of ANNet rose from 0.024 Nm (joint 1) and 0.023 Nm (joint 2) without noise to 0.28 and 0.13 Nm at 1% and to 7.01 and 3.18 Nm at 25%, and it stayed within 1.4% of the analytical median at every level (*p*\* = 0.99 for all ten comparisons; ANNet higher in 45 to 50 of 50 trajectories, paired Wilcoxon signed-rank *p* ≤ 6.71 × 10^−8^). ANNet therefore transmits input noise as the exact equations of motion do, and the learned energy adds at most 1.4% to the median error.

**Fig. 7.**
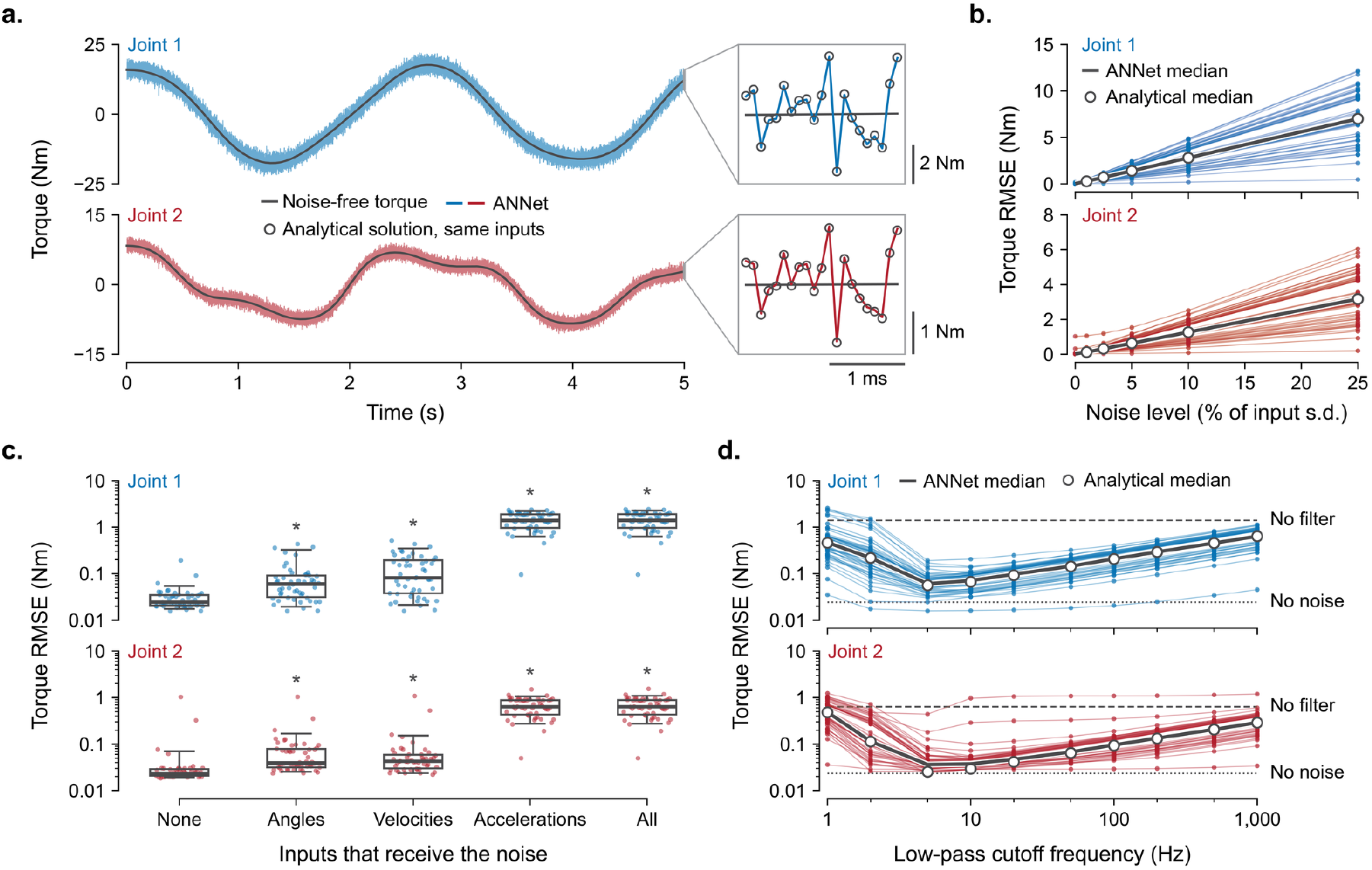
ANNet reproduces the sensitivity of analytical inverse dynamics to measurement noise. **a**, Torques of one trajectory with 5% noise on all inputs. Colored traces, ANNet; black lines, noise-free torques. Frames magnify the final 2 ms, marked at the end of each trace, where the ANNet trace passes through every analytical value (scale bars, 1 ms; 2 Nm for joint 1, 1 Nm for joint 2). **b**, Root-mean-square error (RMSE) of the torques, relative to the noise-free torques, against the noise level. Thin lines join the ANNet errors (dots) of each trajectory (*n* = 50; one noise realization each), heavy lines join ANNet medians and circles mark analytical medians. One-way ANOVA of ANNet against analytical errors, or Kruskal–Wallis tests for joint 2 at 1 and 2.5%, gave *p*\* = 0.99 (Benjamini–Hochberg-adjusted *p*) in all ten joint-by-level comparisons. Two-sided Wilcoxon signed-rank tests paired by trajectory gave *p* between 1.78 × 10^−15^ and 6.71 × 10^−8^. **c**, ANNet RMSE without noise (None) and with 5% noise on the angles, velocities or accelerations alone, or on all inputs. Boxes, median and quartiles; whiskers, 5th and 95th percentiles; dots, trajectories (*n* = 50 per condition); asterisks, *p*\* < 0.05 against None (Kruskal–Wallis tests, 40 comparisons). **d**, RMSE with 5% noise on all inputs after zero-phase low-pass filtering, against the cutoff; symbols as in **b**. Dashed and dotted lines, ANNet medians without filtering and without noise. Filtering at 5 Hz lowered the error of all 50 trajectories (two-sided Wilcoxon signed-rank test, *p* = 1.78 × 10^−15^ for each joint). Joint 1 (blue) is above joint 2 (red); open circles, analytical inverse dynamics on the same inputs. Noise was zero-mean Gaussian, independent across samples and inputs (joint angles, velocities, accelerations), with a standard deviation (s.d.) in percent of the s.d. of each input over the trajectory.

We next asked by which law the torque error grows with input noise and compared the measured errors with first-order propagation of the noise through the inverse dynamics (Fig. 7b). This propagation predicts RMSE = *kp* for an exact model and *RMSE* = [*RMSE*(0)^2^ + (*kp*)^2^]^1/2^ for a model with noise-free error RMSE(0), where *p* is the noise level and the gain *k* follows from the Jacobian of the inverse dynamics without any fitted parameter. The analytical errors followed the proportional law within 0.8% in every trajectory, with median gains of 0.280 Nm (joint 1) and 0.127 Nm (joint 2) per percent of noise and a fitted power-law exponent of 1.00 (range 0.9993 to 1.0001); the ANNet errors followed the quadrature law with a median deviation of 0.3% (largest 7.8%), and the gain computed from the Jacobian of ANNet matched the analytical gain (median ratio 1.0003 for joint 1 and 1.0007 for joint 2). Torque error therefore grows linearly with input noise at a rate fixed by the sensitivity of the dynamics, and ANNet reproduces that sensitivity.

To identify which measurement sets the torque error, we applied the noise to one group of inputs at a time (Fig. 7c). The angles, the velocities or the accelerations alone received the noise at each of the five levels, and the other inputs remained exact. At 5% noise, noise on any one group or on all inputs raised the error above its noise-free value (*p*\* ≤ 1.81 × 10^−7^ for the eight comparisons of joint and condition; error higher in 48 to 50 of 50 trajectories, paired Wilcoxon signed-rank *p* ≤ 8.89 × 10^−15^), but by unequal amounts. Angle or velocity noise alone raised the median 1.7-fold to 3.3-fold, to 0.04 to 0.08 Nm, whereas acceleration noise alone raised it 58-fold (joint 1) and 27-fold (joint 2), to 1.40 and 0.63 Nm, and reproduced more than 99.7% of the median error under noise on all inputs at every level. At equal relative noise, the acceleration signal therefore sets the accuracy of the torque estimate, consistent with the direct transmission of acceleration to torque through the inertia matrix.

Because low-pass filtering is the standard processing of measured kinematics, we filtered the noisy inputs before they entered the models and tested whether this procedure recovers the accuracy of torque prediction (Fig. 7d). Each input passed through a zero-phase fourth-order Butterworth filter with a cutoff between 1 Hz and 1 kHz, and both models received the same filtered inputs. At 5% noise, a 5-Hz cutoff lowered the median ANNet error 23-fold, from 1.40 to 0.060 Nm (joint 1), and 18-fold, from 0.64 to 0.036 Nm (joint 2), against 0.056 and 0.025 Nm for the analytical model; the error fell in all 50 trajectories (paired Wilcoxon signed-rank *p* = 1.78 × 10^−15^), and the filtered medians lay within 2.5 times the noise-free values. Standard low-pass filtering therefore suppresses the noise-driven error of ANNet as it does that of the analytical model.

We then examined what sets the cutoff by comparing the error after filtering with a first-order prediction at every cutoff (Fig. 7d). The prediction combines the error that the filter alone causes on noise-free inputs with the unfiltered error scaled by the fraction of the white-noise standard deviation that the filter transmits. Above 20 Hz the error grew with the square root of the cutoff (log-log slope 0.50 for the analytical model), and the median analytical error stayed within 1.3% of the prediction at every cutoff and noise level; below 5 Hz the filter distorted the movement itself and alone caused median errors of 0.46 and 0.48 Nm at 1 Hz. The cutoff therefore trades transmitted noise against distortion of the movement, as in conventional inverse dynamics, and the transmitted noise scales with the square root of the ratio of cutoff to sampling rate.

### The learned energy preserves the noise tolerance of the forward solver

To test whether the learned energy amplifies noise in the generalized forces beyond the amplification intrinsic to the forward solver, we compared the solver built on ANNet with the same solver built on the analytical Gibbs–Appell energy under the same noise on the gravitational torques (Fig. 8). We added Gaussian noise with a standard deviation of 1, 2.5, 5 or 10% of the standard deviation of each torque at every step of the 50 closed-loop simulations, started every simulation from its exact initial state and scored both solvers against the noise-free reference. The median errors of the two solvers stayed within 13% of each other for angle, velocity and acceleration without noise and at every noise level (ratio of medians 0.90 to 1.13; *p*\* ≥ 0.39 for all 30 comparisons), and at 10% noise ANNet gave the lower error in 32 to 34 of 50 trajectories (paired Wilcoxon signed-rank *p*\* ≤ 0.034). The solver with the learned energy therefore reproduces the noise sensitivity of the solver with the analytical energy to within 13%.

**Fig. 8.**
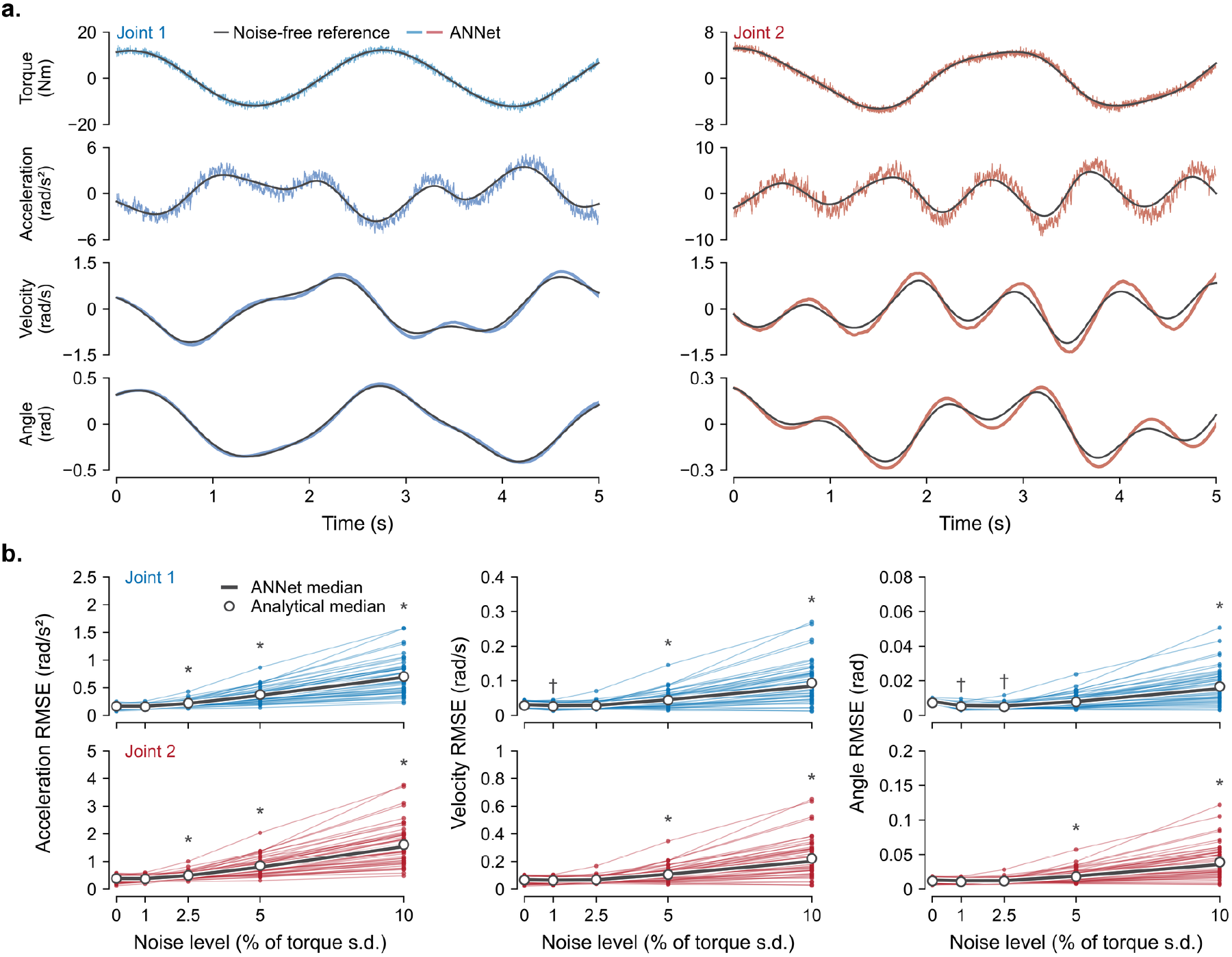
The learned energy preserves the tolerance of the forward solver to noise. **a**, Closed-loop forward simulation of one trajectory with 10% noise on the gravitational torques, joint 1 (blue) on the left and joint 2 (red) on the right. Rows show the gravitational torque that entered the solver and the acceleration, velocity and angle that the solver returned. Colored traces, solver with the learned energy (ANNet); black lines, noise-free fourth-order Runge–Kutta reference. The trajectory is the one whose ANNet errors at 10% noise lay closest to the medians. **b**, Root-mean-square error (RMSE) of acceleration, velocity and angle, relative to the noise-free reference, against the noise level, joint 1 above joint 2. Thin lines join the ANNet errors (dots) of each trajectory (*n* = 50; one noise realization each), heavy lines join ANNet medians and circles mark the medians of the same solver with the analytical Appell energy under the same noise. Asterisks mark the noise levels at which ANNet errors are higher than without noise, and daggers the levels at which they are lower (*p*\* < 0.05, where *p*\* is the Benjamini–Hochberg-adjusted *p* across the 24 comparisons of quantity, joint and noise level; Kruskal–Wallis tests, or one-way ANOVA at the 1% noise level). Unmarked levels have *p*\* ≥ 0.05. Kruskal–Wallis tests of ANNet against analytical errors gave *p*\* ≥ 0.39 in all 30 comparisons of quantity, joint and level. Two-sided Wilcoxon signed-rank tests paired by trajectory gave *p*\* ≤ 0.034 at 10% noise, where ANNet had the lower error in 32 to 34 of 50 trajectories, and *p*\* ≥ 0.20 without noise and at the other levels. Noise was zero-mean Gaussian, independent across joints and solver steps (2 ms), with a standard deviation (s.d.) in percent of the s.d. of each torque over the noise-free trajectory. Every simulation started from the exact initial angles and velocities.

We next compared the errors of the ANNet solver at each noise level with its noise-free errors to establish how much torque noise the forward simulation tolerates (Fig. 8b). Every trajectory entered every condition with the same noise realization and the same candidate accelerations, so that the conditions differed only in noise amplitude. Noise of 1% left the median acceleration error at 1.00 and 0.99 times its noise-free value (*p*\* ≥ 0.44), noise of 2.5% raised it 1.3-fold in both joints (*p*\* ≤ 1.70 × 10^−6^), and noise of 10% raised the median error 4.1-fold and 4.0-fold in acceleration (to 0.69 and 1.54 rad/s^2^ in joints 1 and 2), 2.7-fold and 2.9-fold in velocity (to 0.084 and 0.20 rad/s) and 1.9-fold and 2.7-fold in angle (to 0.015 and 0.036 rad; *p*\* ≤ 2.75 × 10^−8^ for the six comparisons at 10%). At 1 and 2.5% the angle error of joint 1 fell to 0.71 and 0.68 times its noise-free value (0.71 and 0.67 times with the analytical energy), and at 1% its velocity error fell to 0.91 times (*p*\* ≤ 0.012). The relative rise of the error therefore decreased from acceleration to velocity to angle, and the median angle and velocity errors stayed at or below their noise-free values for noise of 1 and 2.5%.

### Inverse-dynamics learning transfers to forward dynamics

To test whether learning of the inverse dynamics transfers to the forward dynamics, we adapted the network to perturbed pendulums by training on torques alone (Fig. 9a,c). We changed the masses, the lengths or both, trained the pretrained network on the torques of each perturbed pendulum for 300 epochs, and ran the forward solver on that pendulum, with its gravitational torques, before and after the adaptation. The adaptation lowered the median torque error 16.8-to 46.8-fold and the median angle error of the forward simulation 4.8- to 11.6-fold, to between 9.6 × 10^−3^ and 13.6 × 10^−3^ rad (joint 1) and between 13.6 × 10^−3^ and 32.9 × 10^−3^ rad (joint 2); the errors of angle, velocity and acceleration fell in 48 to 50 of 50 trajectories (*p*\* ≤ 3.06 × 10^−14^, 18 comparisons). Learning acquired through the inverse dynamics therefore transfers to the forward dynamics without forward-specific training.

**Fig. 9.**
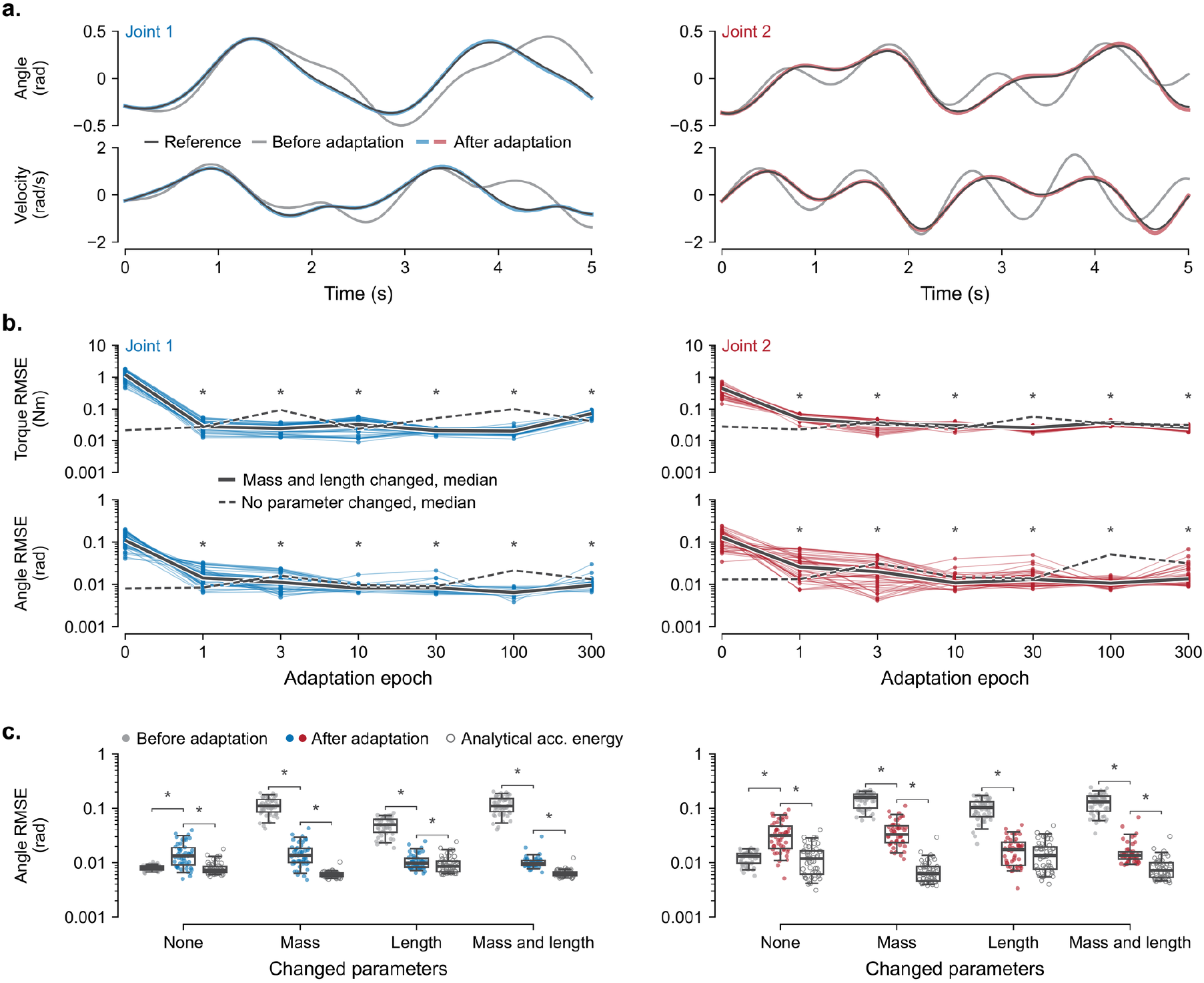
Learning of the inverse dynamics transfers to the forward dynamics without forward-specific training. **a**, Forward simulation of one trajectory (errors closest to the medians) of the pendulum with changed masses and link lengths, joint 1 (blue) on the left and joint 2 (red) on the right. Black lines, fourth-order Runge–Kutta reference; grey and colored lines, forward solver with the network (ANNet) before and after adaptation. The adaptation trained the network for 300 epochs on the torques of the perturbed pendulum alone (masses of 1.2 and 0.8 kg and link lengths of 0.9 and 1.1 m, in place of 1 kg and 1 m). **b**, Root-mean-square error (RMSE) of the torque, the trained quantity, and of the angle of the forward simulation, which never entered the training, at the stored epochs of that adaptation (epoch 0, before adaptation). Thin lines join the errors (dots) of each trajectory (*n* = 50), heavy lines join medians, and dashed lines join the medians of a control, the unchanged pendulum under the same training. Asterisks mark epochs whose errors differ from those at epoch 0 (*p*\* ≤ 2.4 × 10^−16^, where *p*\* is the Benjamini–Hochberg-adjusted *p* of a Kruskal–Wallis test or, for two normal samples, of a one-way ANOVA). **c**, Angle RMSE of the forward simulation of each pendulum (no parameter, the masses, the link lengths or both changed) before adaptation (grey dots), after adaptation (colored dots) and with the analytical acceleration energy (open circles; acc., acceleration). Boxes, median and quartiles; whiskers, 5th and 95th percentiles; dots and circles, trajectories (*n* = 50 per box). A bracket with an asterisk joins a condition to the adapted network when their errors differ (*p*\* ≤ 0.027); the training lowered the error of the perturbed pendulums and raised that of the control. The comparison without a bracket gave *p*\* = 0.14, and two-sided Wilcoxon signed-rank tests paired by trajectory gave *p*\* ≤ 5.4 × 10^−8^ for the eight comparisons with the analytical energy. The solver received the gravitational torques of the simulated pendulum in every condition.

We then asked how well the forward accuracy recovers and compared the adapted solver with three references (Fig. 9b,c). The references were the analytical solver of the perturbed pendulum, the pretrained solver on the original pendulum, and the control solver, whose network received the same training on the original pendulum. The errors of the adapted solver were 1.1 to 6.2 times those of the analytical solver (paired Wilcoxon signed-rank *p*\* ≤ 5.4 × 10^−8^, 18 comparisons) and 0.74 to 2.5 times those of the pretrained solver. The same training raised the errors of the control solver 1.6- to 2.9-fold above those of the pretrained solver (to a median angle error of 0.0133 and 0.0316 rad). The pretrained network is the checkpoint with the lowest validation loss of its 3,000 training epochs, whereas the control and adapted networks hold the weights of epoch 300 without selection. The errors of the adapted solver were 0.26 to 1.04 times those of the control solver (lower in 12 of the 18 comparisons, *p*\* ≤ 0.011, and *p*\* ≥ 0.17 in the other six; with the paired test, lower in 15 and higher in one). Training on torques thus brings the forward error on the perturbed pendulums to at most 1.04 times the error that the same training leaves on the original pendulum.

To establish when the forward error falls during the adaptation, we simulated every pendulum with the network stored after 1, 3, 10, 30, 100 and 300 epochs (Fig. 9b). Each stored network entered the forward solver with the solver configuration held fixed. One epoch lowered the median torque error 9.0- to 43.5-fold and the median angle error 3.2- to 7.7-fold, and the forward errors stayed below those before adaptation at every stored epoch (43 to 50 of 50 trajectories; *p*\* ≤ 4.22 × 10^−14^, 108 comparisons). From epoch 10 the median angle error ranged from 0.0065 to 0.0136 rad (joint 1) and from 0.0104 to 0.0329 rad (joint 2), at or below the largest median error of the control solver at its stored epochs (0.0216 and 0.0512 rad). The forward error falls mostly within the first epoch of training on torques.

Finally, we measured the error that a forward model retains when the training on torques does not reach it, scoring the model without any update against each perturbed pendulum. This model was the pretrained solver with the gravitational torques of the original pendulum, simulated from the same 50 initial states. Its median angle errors were 0.099 rad (joint 1) and 0.21 rad (joint 2) on each of the two pendulums with changed masses and 0.017 and 0.025 rad on the pendulum with changed lengths alone, 1.4 to 15.3 times those of the adapted solver (1.3 to 21.6 times over angle, velocity and acceleration; *p*\* ≤ 0.0049, 18 comparisons). A forward model that the torque data do not reach keeps this error, whereas in ANNet, supplied with the gravitational torques of the changed pendulum, the shared acceleration energy carries the update to the forward dynamics.

## Discussion

This study shows that the inverse and forward dynamics of a mechanical system can be computed from a single learned representation and do not have to be learned as two separate models. A neural network that was trained only on inverse dynamics learned the Appell acceleration energy of a simulated double pendulum. This one function was sufficient to compute the joint torques required for a given movement and, when combined with the applied forces, to predict the movement that those forces produce. The motivation for this work came from sensorimotor control literature, where dynamics models are used in the inverse direction to compute the commands needed for a movement and in the forward direction to predict the consequences of those commands^30,63^. Existing hypotheses propose that the nervous system maintains internal models of both kinds, which are commonly described as paired but separately parameterized modules^31^ and which must adapt whenever the motor apparatus^64^ or the environment^65^ changes. However, it remains unclear whether prediction and control rely on separate representations or on a shared one. Although our simulations cannot reveal how the nervous system is organized, they indicate that this separation is not imposed by the mechanics of the body. Classical mechanics already provides a single function from which both computations can be derived, and our results show that a neural network can learn this function from torque data alone.

The reason that one network can serve both purposes lies in the Gibbs–Appell formulation of mechanics. In this formulation, the acceleration energy is a scalar function whose derivative with respect to acceleration gives the inertial and velocity-dependent forces. Gauss’s principle of least constraint then identifies the true acceleration as the one that minimizes an objective built from this function and the applied forces. Inverse dynamics therefore corresponds to differentiating the learned energy, whereas forward dynamics corresponds to minimizing an objective that contains it, so that a second network is unnecessary. Our work hence extends a family of methods that embed physical structure in neural networks instead of treating dynamics as an unconstrained regression problem. Some of these methods add physical knowledge to the training objective^66^. Others build the structure into the network itself, for example by learning an inertia matrix that is positive definite by design^67^, a Hamiltonian or a Lagrangian^68–70^, or a port-Hamiltonian system^71^. Like the energy-based networks among them, ANNet learns a scalar quantity and obtains forces from its derivatives, and its loss is related to Sobolev training^54^, although ANNet is supervised through the derivative alone. ANNet differs from these networks mainly in how the two computations are obtained, because the torque follows from a single derivative with respect to acceleration and the acceleration follows from an optimization. Consistent with this literature, imposing physical structure improved accuracy, as ANNet predicted joint torques with a lower median error than an otherwise identical network that was trained to output the torques directly. A plausible explanation is that the energy formulation enforces relationships that a direct regressor would have to discover from data. For example, because every torque is a partial derivative of the same function, the inertial coupling between the two joints is symmetric by construction. The behavior of ANNet during training also agreed in part with earlier reports on physics-informed networks. In those reports, accuracy was limited more by the difficulty of optimization than by network capacity^72^, and additional width and depth beyond a certain point brought little benefit^73,74^. In our hyperparameter search, performance depended mostly on the width of the first hidden layer and on the learning rate. The choice of activation function, which has been reported to influence both the achievable error^75–77^ and the speed of convergence^78^, had only a minor effect.

A further indication that ANNet learned the structure of the dynamics is that it responded to noise in nearly the same way as the analytical equations of motion, although it was trained on exact (“clean”) data. Inverse dynamics is known to amplify uncertainties in its inputs^79^, and the torques predicted by ANNet showed nearly the same sensitivity to measurement noise as torques computed from the analytical equations, with most of the error arising from noise in the accelerations. Accordingly, the low-pass filtering that is routinely applied in conventional inverse dynamics was also effective for ANNet. In the forward direction, white noise in the applied torques degraded the simulations about as much with the learned energy as with the analytically computed energy. When we replaced the learned energy with the analytically computed one, the accuracy of the forward simulations remained nearly unchanged within the sampled range of states, which indicates that the remaining error originated in the numerical solver and the integration scheme and not in the network. Keeping the learned equations of motion separate from numerical integration also made forward dynamics more stable in our earlier work^80^. Because both computations draw on the same learned energy, learning in the inverse direction should also reach the forward direction, and our simulations confirmed this expectation. When the masses and link lengths of the pendulum were changed and the network was retrained using only the torque error, the forward simulations of the changed pendulum improved as well, even though they never entered the training and only the gravitational torques of the changed pendulum were supplied to the solver. In schemes that give the forward and the inverse model separate parameters and separate training signals^31^, by contrast, each model has to learn such a change from its own training signal.

Several limitations should be considered when interpreting these findings. The most important one is that our evidence comes from a single low-dimensional system, which was simulated from analytical equations and moved passively under gravity. The planar double pendulum is an established model of the interaction torques that arise during reaching^81,82^ and of the passive mechanics of walking^83–85^, but it lacks actuation, contact, compliance and sensorimotor delays. The principle is not tied to this system, because the Gibbs–Appell formulation holds for constrained mechanical systems in general, but whether the acceleration energy of more complex systems can be learned with comparable accuracy remains to be shown. A second limitation is that ANNet was accurate only within the range of states on which it had been trained, and the largest torque errors occurred where test trajectories left that range. The forward simulations remained inside the sampled range, however, so this restriction does not affect the conclusion that the learned energy supports forward dynamics. A third limitation is that robustness was assessed only with white noise, while gravity was supplied analytically and the training torques were exact. These choices isolated the inertial terms that our hypothesis concerns. Because ANNet responded to noise in nearly the same way as the analytical equations, the methods that are already used to handle measurement errors in inverse dynamics are likely to remain applicable, although this needs to be confirmed with recorded data. The comparison with direct regression also has a caveat, because the regressor used the architecture that had been optimized for ANNet and might perform better with an architecture of its own. Even in that case, however, a regressor would provide only the inverse computation, whereas the learned energy provides both. The network itself is not constrained to be quadratic in the accelerations, so a single minimum of the objective is not guaranteed, although the solver was about equally accurate with the learned and the analytical energy. Finally, the forward solver relies on a random search over candidate accelerations and on a penalty that discourages abrupt changes in acceleration. This design is expected to become costly as the number of joints increases, it has not been tested with sudden perturbations, and although the average update met the real-time limit, complete simulations were slower than numerical integration of the analytical equations. These shortcomings concern the solver that we used and not the formulation itself, and the solver can be retuned or replaced without retraining the network.

Looking ahead, the most direct next step is to apply this approach to articulated systems with more degrees of freedom and to train the network on measured torques instead of simulated ones. Classical system identification specifies the form of the rigid-body equations in advance and estimates their parameters from motion and torque data^86–88^, which are typically collected along exciting trajectories designed for this purpose^89–91^. Data-driven alternatives learn inverse dynamics directly from examples^92^ and are already used to control robotic manipulators^93,94^, but they provide only one of the two computations. Learning the acceleration energy from the same kind of data could, in principle, yield a single model that supports both. Forces that depend only on position and velocity, such as gravity and friction, could be learned by the same network, because they enter the energy as terms that are linear in acceleration. Contact forces and muscle forces that depend on activation dynamics would have to be supplied separately. Torque-controlled robots are well suited for an initial test of this idea, because their joint torques are measured and their accelerations can be varied independently of the state. Wearable robots present a harder case, since only the torque of the device is measured and the contribution of the user has to be estimated independently. Such a model would complement physics engines, which remain the method of choice when a complete rigid-body description is available and many fast simulations are required^95–97^, and it would be most useful for systems whose equations are difficult to write down but whose motion and torques can be recorded. For systems with many joints, the random search in the forward solver could be replaced by a single-step solution that starts from the acceleration of the previous time step, because the analytical acceleration energy is quadratic in the accelerations, although we have not yet tested this possibility. Our results also point to a further exploration of sensorimotor control strategies. If prediction and control rely on a shared representation of limb dynamics, then adaptation acquired in one of them should transfer to the other, whereas models with separate parameters and separate training signals predict no such transfer. Observing this transfer during human motor learning would be consistent with a shared representation, and its absence would argue against one. More broadly, our findings indicate that predicting a movement and producing it do not have to be treated as separate modeling problems, because a single function that is grounded in classical mechanics and learned from data can serve both.

## Conclusions

Learning the Appell acceleration energy of a simulated double pendulum from joint torques allowed a single neural network to solve both inverse dynamics, by computing the torques required for a movement, and forward dynamics, by predicting the motion that applied forces produce. This demonstrates that inverse and forward dynamics can be learned as one model, without training a separate network for each. Ultimately, this approach paves the way for robots that learn their own dynamics from kinematic and kinetic data and use one model for both inverse and forward dynamics.

## Data Availability

All data generated in this study are deposited in the figshare repository^98^ and are mirrored in the GitHub repository https://github.com/SerhiiBahdas/ms-annet. The processed data required to reproduce the manuscript figures are provided in SUBMISSION/FIGURES/DATA AND SCRIPTS/ as figure-specific archives and notebooks. The saved simulation outputs, model checkpoints, timing measurements, training histories, normalization statistics, and hyperparameter-search results are provided in SIMULATIONS/. The source data for Figs. 2 to 9 are in Supplementary Data 1 to 8, and the source data for Supplementary Figs. 1 to 3 are in Supplementary Data 9 to 11. No external empirical dataset was used; the data were generated from the double-pendulum simulations and models described in the manuscript.

## Code Availability

All code required to reproduce the analyses and figures in this manuscript is available in the GitHub repository https://github.com/SerhiiBahdas/ms-annet under the MIT License, without restrictions on access, and the version used in this study is available under a figshare repository^98^. The repository includes run instructions in README.md, figure-generation notebooks in SUBMISSION/FIGURES/DATA AND SCRIPTS/, and the full simulation and analysis workflows in SIMULATIONS/MAIN_RUN_ME_NEW.ipynb and SIMULATIONS/HYPERPARAMETERS_RUN_ME.ipynb. The analyses ran in Python 3.13.11 with PyTorch 2.11.0, NumPy 2.3.5, SciPy 1.16.3, statsmodels 0.14.6, SymPy 1.14.0, Optuna 4.8.0, scikit-learn 1.8.0, pandas 2.3.3 and Matplotlib 3.10.8. The file requirements.txt pins these versions, and the notebooks set the random seeds and all analysis parameters.

## Supplementary information

### Appellian Neural Networks

Serhii Bahdasariants, Lauren Parola, Kriti Kacker, Ariel K. Feldman, Zachary Zdobinski, Inseung Kang, Douglas J. Weber

### Forward dynamics equations

Let **q** = [*q*_1_, *q*_2_]^*T*^ denote joint angles and 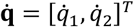 denote angular velocities. From the Gibbs–Appell formulation, the partial derivative of the Appell acceleration energy with respect to generalized acceleration can be written as:

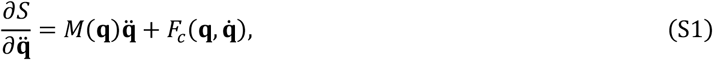

where *M*(**q**) is the configuration-dependent mass matrix and 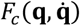 contains velocity-dependent inertial terms. Setting this expression equal to the negative gradient of the potential energy gives:

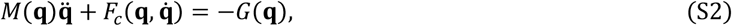

where *G*(**q**) = *∂V*/*∂* **q**. Thus, the ground-truth forward dynamics used for numerical integration were obtained as:

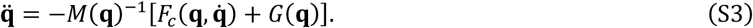

For the two-link system, the matrix and vectors were:

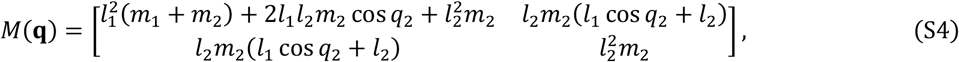

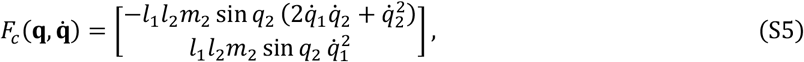

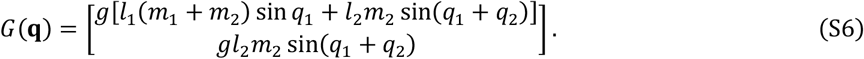

Equivalently, defining 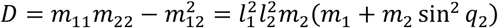, the joint accelerations were:

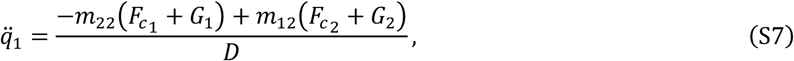

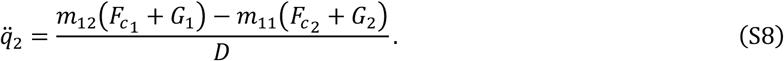

These accelerations were then reformulated as the first-order system:

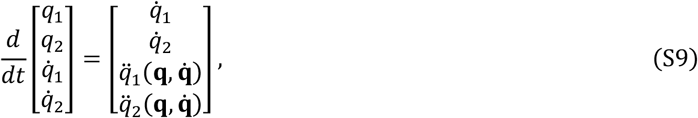

which was integrated using the fixed-step fourth-order Runge–Kutta scheme.

### Supplementary Figures and Legends

**Supplementary Fig. 1.**
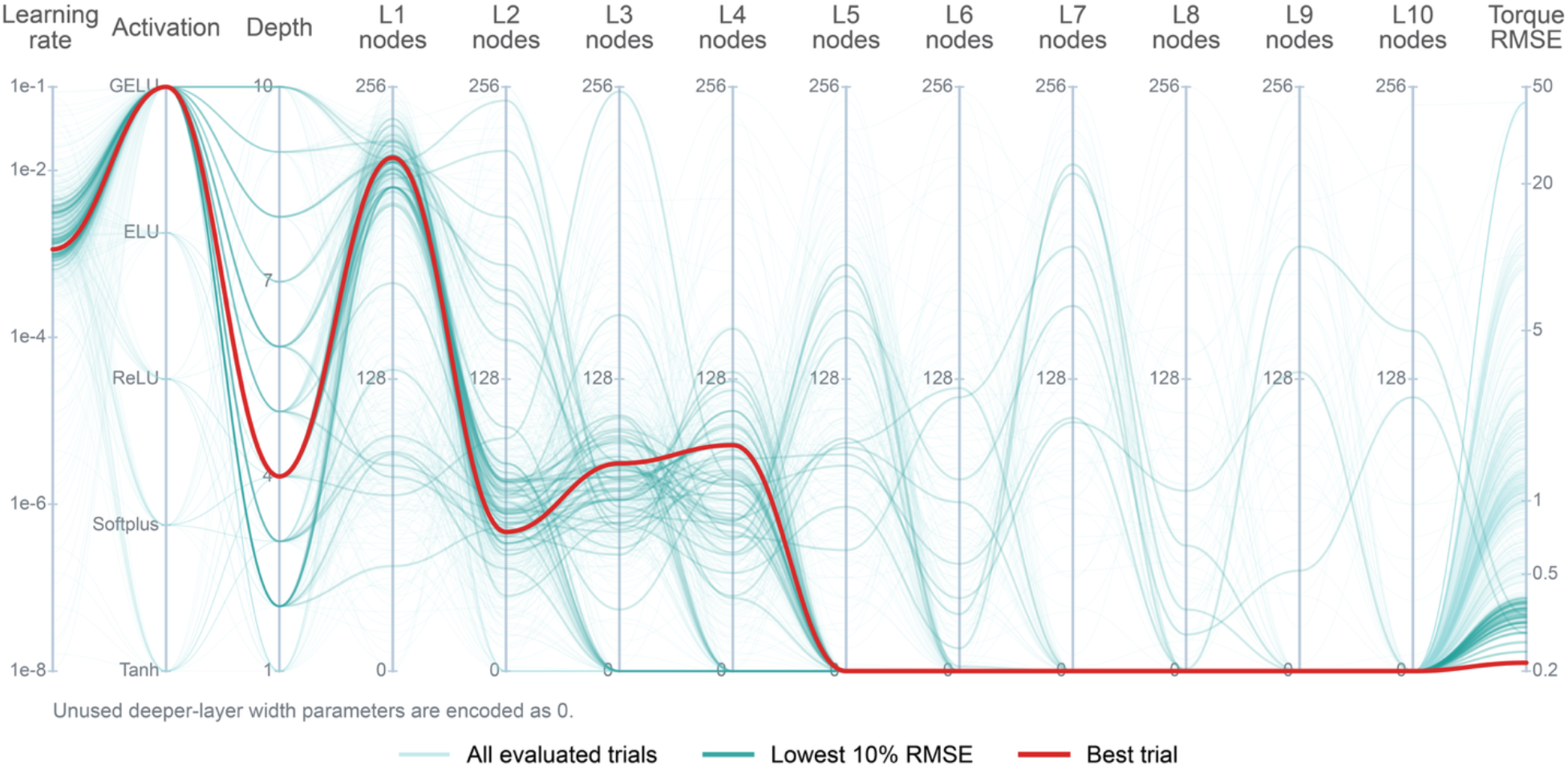
Parallel-coordinate representation of hyperparameter optimization. Each polyline represents one Optuna hyperparameter trial evaluated during the architecture search. Axes show the learning rate, activation function, network depth, hidden-layer widths, and validation torque RMSE. Pale cyan lines show all evaluated trials (n = 500), teal lines indicate the lowest 10% of trials by RMSE (n = 50), and the red line marks the best-performing configuration used for subsequent analyses. Unused deeper-layer width parameters in shallower networks are encoded as 0 to display variable-depth architectures on a common axis set.

**Supplementary Fig. 2.**
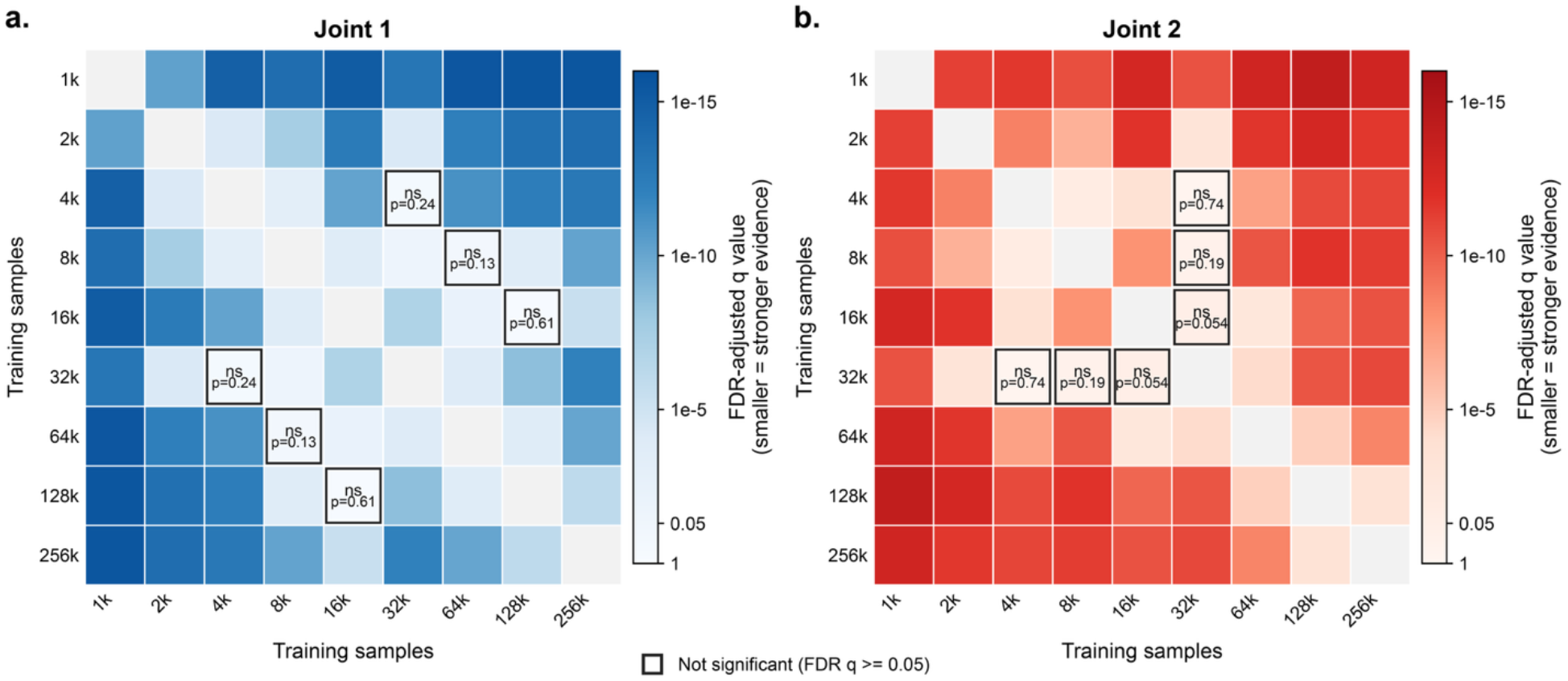
Pairwise statistical comparison of inverse-dynamics accuracy across training-set sizes. **a**,**b**, Matrix representation of pairwise comparisons of trajectory-wise torque RMSE distributions (n = 50 test trajectories per training-set size) for joint 1 (**a**) and joint 2 (**b**) across training-set sizes. Rows and columns denote the dataset sizes being compared. Pairwise differences were evaluated using two-sided Mann–Whitney U tests with Benjamini–Hochberg false discovery rate correction. Cell color reports the FDR-adjusted p value, with darker blue for joint 1 and darker red for joint 2 indicating stronger evidence that the corresponding RMSE distributions differ. Gray diagonal cells represent self-comparisons. Outlined cells labeled ns denote comparisons that were not significant after FDR correction (p ≥ 0.05).

**Supplementary Fig. 3.**
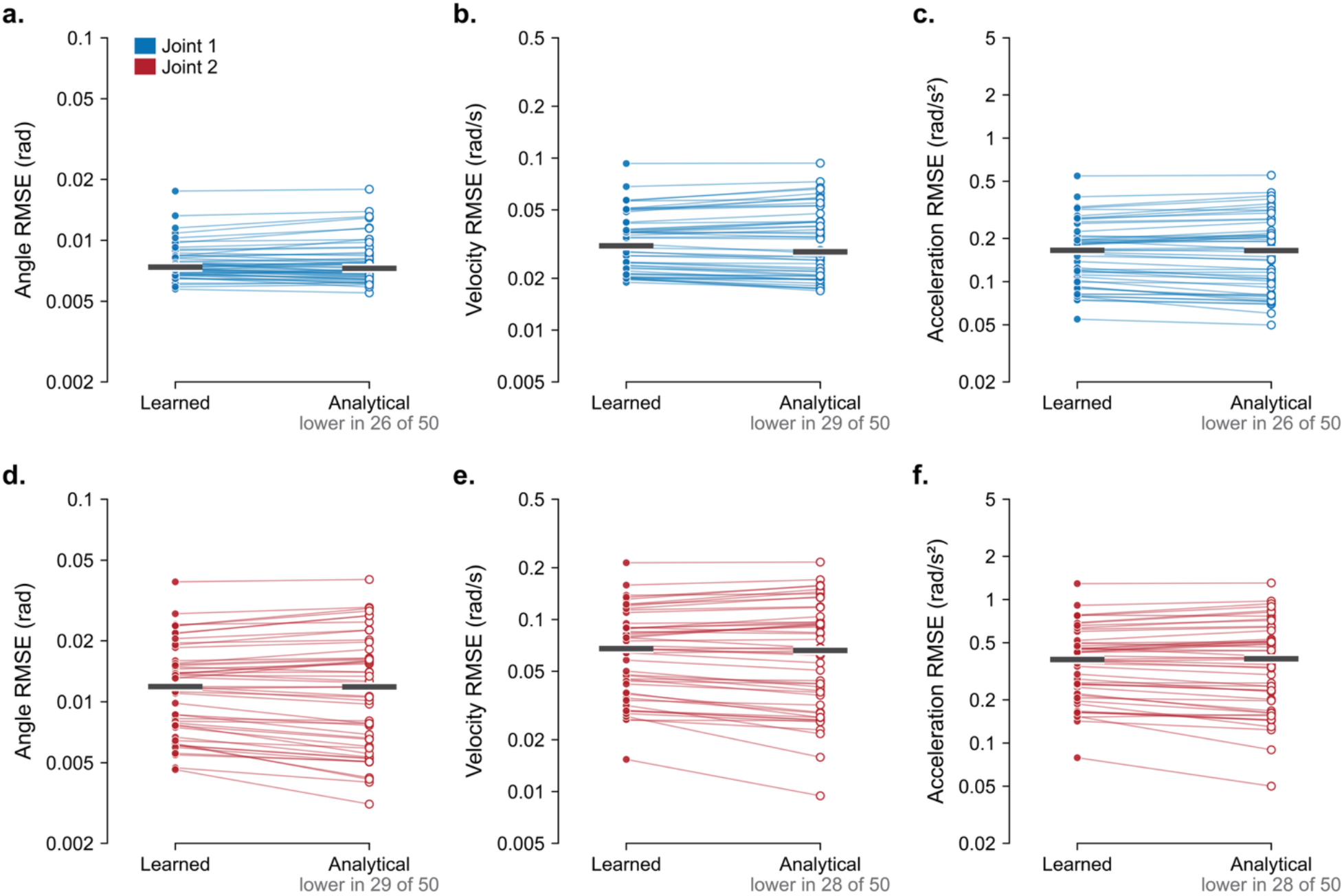
Learned and analytical acceleration energies give median closed-loop simulation errors within 8% of each other. **a–f**, Root-mean-square error (RMSE) of joint angle (**a, d**), velocity (**b, e**) and acceleration (**c, f**) of the closed-loop forward simulation for joint 1 (**a–c**, blue) and joint 2 (**d–f**, red), on logarithmic axes. Filled circles, solver with the learned acceleration energy (Learned; the errors of Fig. 6b); open circles, the same solver with the analytical Appell acceleration energy substituted for the learned energy (Analytical). Each line joins the two errors of one trajectory (*n* = 50), heavy bars mark medians, and the count beneath each panel gives the trajectories in which the analytical energy had the lower error. Both simulations used the same solver configuration (16 candidate accelerations within ±15 rad/s^2^, 10 Adam refinement steps, *λ* = 0.1), semi-implicit Euler steps of 2 ms, initial states and fourth-order Runge–Kutta reference trajectories of 5 s. The medians with the analytical energy were 0.92 to 1.01 times those with the learned energy (Kruskal–Wallis tests, *p*\* = 0.94 for the six comparisons, where *p*\* is the Benjamini–Hochberg-adjusted *p*; two-sided Wilcoxon signed-rank tests paired by trajectory, *p* between 0.41 and 0.98).

